# Ribosome Processing Factor-2 Interacts with RPL10A to Regulate Selective Translation during Plant Immunity and Drought Stress

**DOI:** 10.64898/2026.03.12.711238

**Authors:** Shobhna Yadav, Kesia Mathew, Supriya Singh, Ankit Biswas, Sanjay Deshpande, Chanchal Kumari, Saritha K Reddy, Keri Wang, Tushar K Maiti, Kirankumar S. Mysore, Ramu S. Vemanna

## Abstract

Processing of ribosomal RNA (rRNA) is essential for ribosome biogenesis, translation, plant development, and stress adaptation. Ribosome processing factor-2 (RPF2), which plays a role in the later stages of rRNA maturation, interacts with ribosomal protein L10A (RPL10A). *RPF2* overexpression in Arabidopsis and *Nicotiana benthamiana* showed enhanced plant growth and trichome development due to increased gibberellic acid (GA) levels. Conversely, *RPF2*-silenced and mutant plants had a dwarf phenotype, reduced stomatal apertures, and decreased glucosinolate accumulation. *RPF2* silenced and mutant plants also showed compromised nonhost disease resistance, whereas *RPF2* overexpression lines exhibited enhanced disease resistance to both host and nonhost pathogens. *RPL10A* and *RPF2* overexpression lines were sensitive to abscisic acid (ABA) and tolerant to drought, which is attributed to their unique roles in translation regulation. Despite having larger stomatal apertures, *RPF2* overexpression plants displayed low pathogen multiplication rates and reduced water loss, indicating independent resistance mechanisms associated with ribosomal functions in translation regulation. Although both RPL10A and RPF2 proteins interact with each other and are involved in translation regulation, proteomic analysis suggests that they regulate the translation of distinct sets of genes during pathogen or drought stress. These findings indicate that RPF2 and RPL10A play independent roles in the regulation of unique protein translation.

## Introduction

In eukaryotic cells, ribosome biogenesis begins with the synthesis of 45S pre-rRNA, which forms a 90S pre-ribosomal particle within the nucleolus. This particle then undergoes several transformations to yield 25S, 5.8S, and 5S rRNAs. In plants, rRNA maturation is a complex multistep process involving various alternative pathways and numerous ribosome biogenesis factors (RBFs), ribosomal proteins (RPs), and small nucleolar RNAs (Henras & Plisson-chastang, 2015; Weis et al., 2015). Initially, 45S rRNA is processed by nucleolin, H (pseudouridine)/ACA box complex proteins, and exo/endo-nucleases, followed by the involvement of Interacting with MPP10 (IMP4) / Biogenesis of Ribosomes in Xenopus (BRIX) family proteins in the later stages of maturation. Arabidopsis contains six RNA-binding factors (RBFs) with BRIX domains, but their molecular function remains unknown (Maekawa et al., 2018). BRIX2/Ribosomal RNA processing factor 2 (RPF2) forms a complex with Regulator of Ribosome Synthesis (RRS1) to bind 5S rRNA, contributing to the formation of the pre-60S ribosomal subunit alongside ribosomal protein L5 (RPL5) and RPL11 (Asano et al., 2015; Khadre et al., 2015; Madru et al., 2015). Arabidopsis RPF2 (ARPF2) interacts with 5S ribosomal RNA (rRNA) or its precursor, as well as with spacer 2 in the precursors of 25S rRNA (Maekawa et al., 2018). Overexpression of *ARPF2* and *ARRS1* in Arabidopsis leads to shorter stems, abnormal leaf shapes, and prolonged reproductive growth (Maekawa et al., 2018). Transient silencing of *ARPF2* in Arabidopsis using *Tobacco rattle virus* (TRV)-mediated virus-induced gene silencing (VIGS) also resulted in similar phenotypes (Choi et al., 2020). Arabidopsis BRX1, BRX1-1, BRX1-2, and BRIX2/RPF2 are implicated in pre-rRNA processing (Weis et al., 2015). Null mutants of Arabidopsis homologs of yeast Ssf1/Ssf2, SNAIL1, and RPF2 are lethal (Hao et al., 2017; Maekawa et al., 2018). Mutations in ribosome biogenesis genes lead to developmental defects, embryo lethality, reduced plant size, altered leaf shape, reduced cell division, decreased fertility, and defects in trichome development (Byrne 2009). Nucleolin, Fibrillarin2, Hot5, Small organ, Ribosomal RNA processing 7, Arabidopsis Pumilio protein (Apum) 24, and other mutants exhibit dwarf, shrivelled phenotypes with defects in the accumulation of 32S, 27S, 18S, and 5.8S rRNA (Hang et al., 2014). Despite their role in plant development, the function of RPF2 in plant stress responses remains unclear.

Previous studies have examined the function of ribosomal proteins in both basal and innate immunity against pathogens. Silencing *RPL10A, B, RPL12*, and *RPL19* in *Nicotiana benthamiana* and Arabidopsis resulted in weakened nonhost disease resistance and increased susceptibility to various bacterial pathogens (Nagaraj et al., 2016; Ramu et al., 2020). RPL10A regulates transcription and translation and is crucial for plant growth, development, and immune responses to pathogens (Ferreyra et al., 2010; Ramos et al., 2020; Ramu et al., 2020). Ribosomal RNA and ribosomal proteins are essential for the fundamental metabolic activities of any cell and influence stress adaptation. In this context, investigating the molecular mechanisms by which rRNA processing factors contribute to plant tolerance to biotic and abiotic stress could offer an avenue for developing climate-resilient crops. In this study, we demonstrated that silencing *NbRPF2* in *N. benthamiana* or mutant *AtRPF2* in Arabidopsis reduced disease resistance to both host and nonhost pathogens. We identified that AtRPF2 interacts with proteins involved in ribosome biogenesis, particularly through its interaction with AtRPL10A. Overexpression of *RPF2* in Arabidopsis and *N. benthamiana* resulted in increased plant biomass, enhanced stress tolerance, and improved protein translation. Similarly, *RPL10A* overexpression boosted disease and drought stress tolerance. RPF2, along with RPL10A, regulates the translation of specific proteins in response to pathogen infection and drought stress.

## Results

### Modulation of *RPF2* expression in *N. benthamiana* affects nonhost disease resistance and plant development

Downregulation of cDNA clone *NbME19A10*, identified in a previous study (Senthil-kumar et al., 2018), in *N. benthamiana* via TRV-based VIGS led to a dwarf and crinkled phenotype (Fig. S1A) and reduced resistance to the nonhost pathogen *Pseudomonas syringae* pv. *tomato* T1 (Fig. S1B). The silenced plants exhibited more than 50% downregulation of RNA corresponding to *NbME19A10* cDNA (Fig. S1C) and failed to trigger a hypersensitive response (HR) when the nonhost pathogen was syringe-infiltrated at a concentration of 1×10^4^ cfu mL^-1^ (Fig. S1D). These plants also showed increased fluorescence accumulation when vacuum-infiltrated with *GFPuv* (Wang et al., 2007) expressing host pathogen *P. syringae* pv. *tabaci* or nonhost pathogen *P. syringae* pv. *tomato* T1 at 1×10^4^ cfu mL^-1^ concentration (Fig. 1A&C). The *NbME19A10*-silenced plants had no significant effect on host pathogen growth (Fig.1B). However, nonhost pathogens multiplication increased ∼ two-fold at 7 dpi compared to the non-silenced control (*TRV2::00*) plants (Fig. 1D). The domain architecture of *NbME19A10* cDNA revealed the BRIX domain, a characteristic feature of the ‘Interacting with MPP10 Mitotic (M)-Phase Phosphoprotein 10 [MPP10] protein 4’ (IMP4) superfamily members, which are associated with ribosomal RNAs and play distinct roles in ribosome biogenesis. Through BLAST search, we showed that *NbME19A10* is a homolog of Arabidopsis *RPF2* (*AtRPF2*). RPF2 contains a small Rho-GTPase domain and is conserved across vertebrates, mammals, and plants (Fig.S2A&B). Phylogenetic analysis using the nearest-neighbour model indicated that RPF2 in monocots and dicots formed a distinct clade (Fig. 1E). Plant RPF2 proteins are distinct from those in fungi and animals, suggesting specific roles.

**Figure 1.**
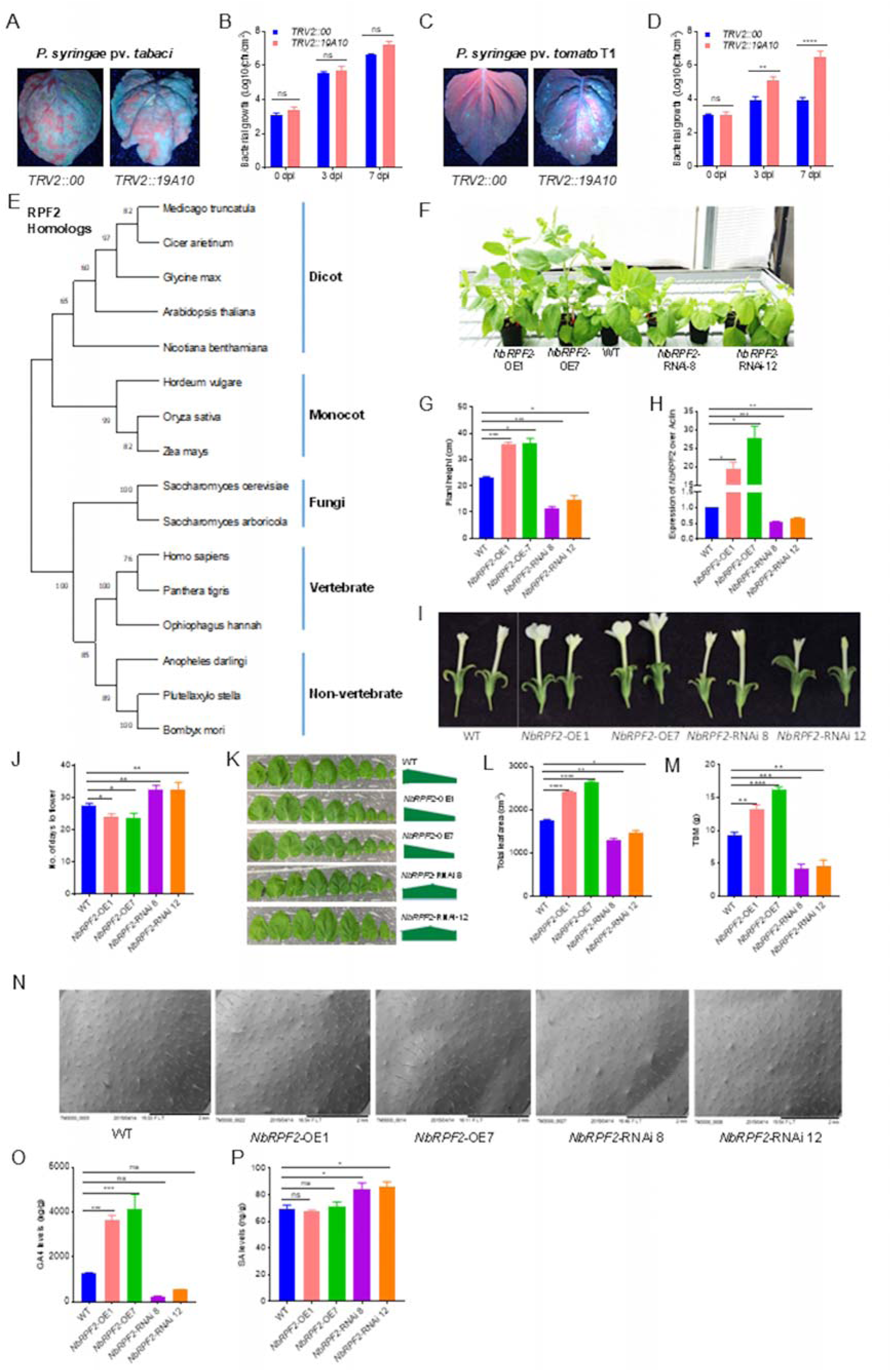
RPF2’s role in nonhost disease resistance and plant development. A. Visualization of *P. syringae* pv. *tabaci* (*pDSK-GFPuv*) in *NbME19A10*-silenced *N. benthamiana* leaves under UV illumination. B. Growth of host-pathogen *P. syringae* pv. *tabaci* at 0, 3, and 7 days after inoculation (dpi). C. Visualization of nonhost pathogen *P. syringae* pv. *tomato* T1 *(pDSK-GFPuv*) in *NbME19A10*-silenced *N. benthamiana* leaves under UV illumination. D. Pathogen growth measurements at 0, 3, and 7 dpi. Four-week-old *N. benthamiana* plants were agro-infiltrated with *TRV::00* or *TRV::19A10*, and three weeks later, *P. syringae* pv. *tabaci* or *P. syringae* pv. *tomato* T1 at 1x10^4^ cfu/ml concentration was vacuum infiltrated, and growth was assessed after 0, 3, and 7 dpi. Photographs were taken at 7 dpi under UV light. The experiments were repeated three times with biological replicates. E. Phylogenetic analysis of AtRPF2 amino acid sequences using 1000 bootstrap values with the MEGA-XI tool. F. Phenotypes of *NbRPF2*-OE and RNAi lines in *N. benthamiana.* G. Plant height after 5 weeks, H. Transcript levels of *NbRPF2* in the leaves of WT, *NbRPF2*-OE and *NbRPF2*-RNAi lines were measured by RT-PCR. The bars represent differential transcript levels compared to WT. Expression of *Actin* gene was used for normalization. I. Flower phenotype from *NbRPF2*-OE, *NbRPF2*-RNAi lines, and WT, J. Days to flowering- minimum 5 plants from each lines were observed for flowering and average is presented., K. Leaf arrangement- all the leaves from on representative plant from the top to bottom and arranged serially and photographed, and L. Total leaf area- all the leaves from three independent plants from each lines was measured using leaf area meter, M. Total dry biomass measured after the harvest of plants. Two independent OE and RNAi lines were used; a minimum of 3 biological replicates were used for all the parameters. N. Scanning electron microscopy images of trichomes from adaxial surfaces (N=20/Plant, Scale-2mm), O. Gibberellic acid (GA4) and P. Salicylic acid (SA) levels in WT, *NbRPF2*-OE and *NbRPF2*-RNAi lines. Three independent lines were evaluated for physiological, morphological, and biochemical characteristics. Error bars represent SE, with two-way ANOVA indicating p*<0.05, p**<0.01, p***<0.001, p****<0.0001.

To investigate the molecular functions of *RPF2, NbRPF2* was stably overexpressed and silenced by RNAi in *N. benthamiana* (Fig. S2C and D). Plants with *NbRPF2* overexpression (*NbRPF2*-OE) exhibited a more vigorous growth pattern than wild-type (WT) plants, whereas *NbRPF2* RNAi plants exhibited reduced growth (Fig.1F). The *NbRPF2*-OE plants were taller, whereas the *NbRPF2* RNAi plants were shorter than the WT (Fig.1G). The severity of the growth phenotype correlated with transcript levels in each line (Fig.1H). Overexpression resulted in early flowering (Fig.1I and J), with no changes in leaf phyllotaxy or shape (Fig.1K), larger leaf area (Fig.1L), increased total biomass (Fig.1M), and displayed spirally arranged leaves (Fig. 1K), allowing all leaves to receive even light, thereby enhancing photosynthesis and increasing biomass. In contrast, the *NbRPF2* RNAi lines showed altered phyllotaxy, with upper leaves that were moderately larger and broader in the middle, shading the smaller basal leaves, reducing light exposure, and resulting in a dwarf phenotype with lower biomass. *NbRPF2*-OE plants also had longer and denser trichomes or leaf hairs than WT and *NbRPF2* RNAi plants (Fig.1N). The *NbRPF2*-OE plants exhibited more than a two-fold increase in GA4 levels, whereas the *NbRPF2* RNAi plants exhibited decreased levels (Fig.1O). *NbRPF2* RNAi plants showed more than a 1.5-fold increase in SA levels, with no significant change observed in *NbRPF2*-OE plants (Fig.1P).

### RPF2 function in plant development and metabolism is conserved in Arabidopsis

To assess RPF2 expression in Arabidopsis in response to different pathogens, publicly available datasets were used. The expression of *AtRPF2* was induced by more than two-fold in response to the host pathogen *P. syringae* pv. *maculicola* and *tomato* (DC3000) at 24 hpi, respectively. However, its expression increased more than four- to six-fold in response to the nonhost pathogen *P. syringae* pv. *tomato* T1, and *Xanthomonas campestris* pv. *vesicatoria* at 24 hpi (Fig. S3). To explore the molecular mechanisms of RPF2, an Arabidopsis homolog (*AtRPF2*) with 66.8% nucleotide and 67.83% amino acid homology to *NbRPF2* was identified (Fig. S4A&B). Arabidopsis *AtRPF2* overexpression (OE) and *AtRPF2* RNAi lines were developed. T-DNA insertion mutants of *AtRPF2* (*Salk_020229* and *Salk_0796070*) were obtained with the T-DNA insertions located in the 3’-UTRs of *AtRPF2* (Fig. S4C). These mutants exhibited reduced *AtRPF2* expression and a slightly dwarf phenotype similar to that of RNAi plants (Fig. 2A and 2B) when compared to Col-0 and *AtRPF2* OE lines. The overexpression lines had longer and more densely distributed trichomes than the RNAi, mutants, and Col-0 plants (Fig. 2C). *AtRPF2*-OE plants accumulated more than twice the levels of GA compared to the RNAi and mutant lines (Fig. 2D). While SA levels in *AtRPF2*-OE lines showed no significant change from Col-0, RNAi and mutant lines exhibited over 2.5-fold higher SA levels than Col-0 (Fig. 2E). Targeted metabolite profiling using methanolic extracts revealed higher glucosinolate levels in *AtRPF2*-OE plants than in Col-0. Glucosinolates such as Indol-3-ylmethyl-glucosinolate, 4-methylsulfinyl-n-butyl-glucosinolate, isorhoifolin (Apigenin-7-O-rutinoside), spiraeoside (quercetin 4’-O-glucoside), m-coumaric acid, and methylsulfinyl-n-heyl-glucosinolates accumulated at significantly higher levels than in Col-0, RNAi, and mutant lines (Fig. 2F). Additionally, long-chain fatty acids, such as linoleic, palmitic, linolenic, and stearic acids, were found at higher concentrations in *AtRPF2*-OE plants. The *AtRPF2*-OE lines displayed a higher stomatal aperture on both adaxial and abaxial leaf surfaces compared to RNAi lines and Col-0 plants (Fig. 2G-I).

**Figure 2.**
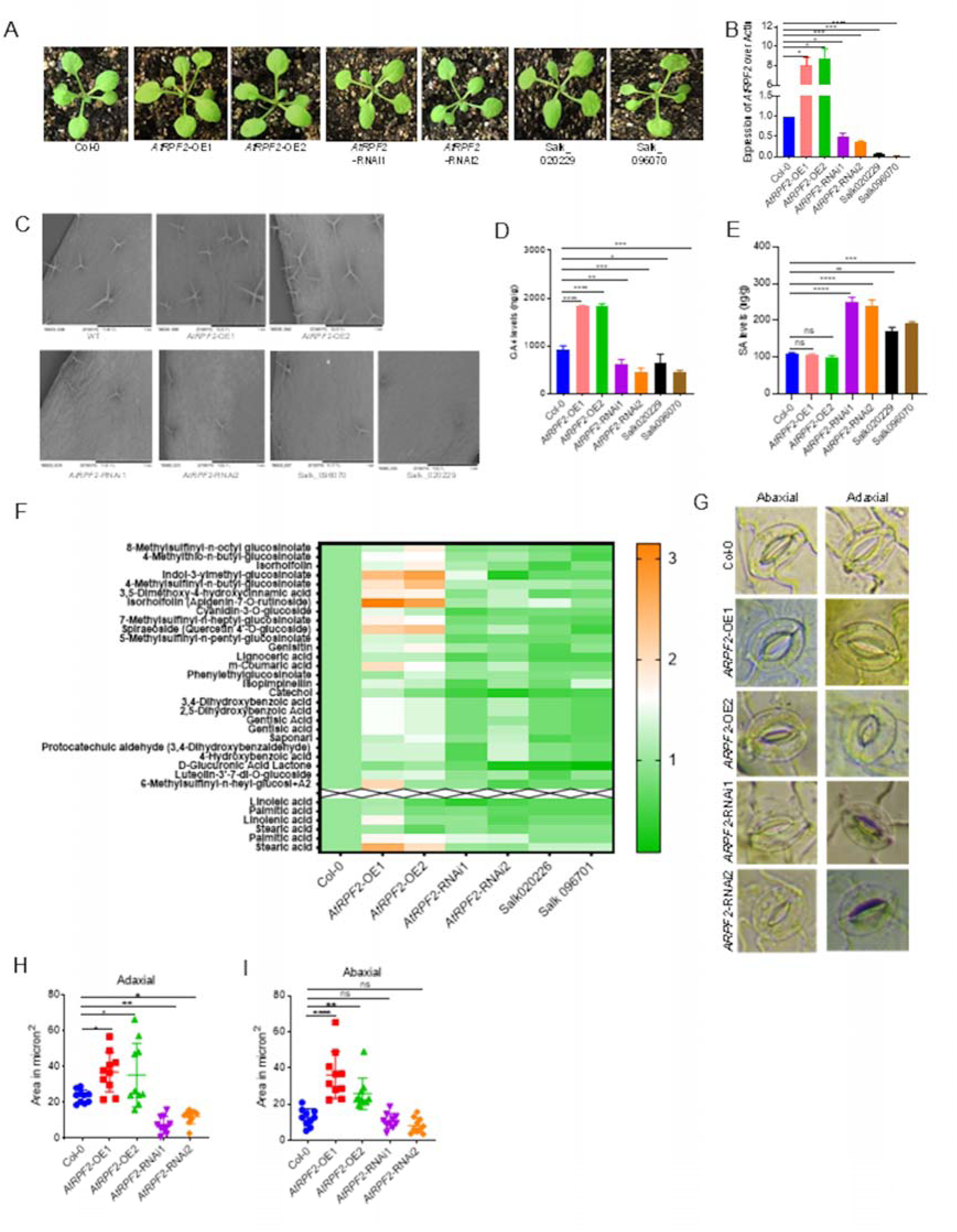
AtRPF2 plays a role in the development of trichomes and the production of secondary metabolites. A. The phenotype of two independent *AtRPF2* overexpression lines (*AtRPF2*-OE1 and *AtRPF2*-OE2), *AtRPF2*-RNAi lines (*AtRPF2*-RNAi1 and *AtRPF2*-RNAi2), and *Atrpf2* mutant lines (Salk_020229 and Salk_096070) after three weeks of germination. B. Transcript levels of *AtRPF2* in OE, RNAi and mutant plants after three weeks of germination. *AtActin* was used for normalization. Three biological replicates from each line were used for total RNA isolation C. Trichome images from adaxial leaf surfaces of 4-week-old *AtRPF2*-OE and RNAi plants were taken using SEM (n=30 from 3 different plants). Phytohormones D. Gibberellic acid (GA4), E. Salicylic acid (SA) levels in four-week-old Arabidopsis plants quantified using HPLC coupled -MS analysis. F. Levels of secondary metabolites and glucosinolates from four-week-old Arabidopsis plants. A non-targeted metabolite analysis was conducted with methanol extracts using GC-MS analysis. The relative values were plotted using GraphPad Prism. G. Photographs depicting stomatal openings. Stomatal area on H. Adaxial and I. Abaxial leaf surfaces (n=20). Epidermal peels from three-week-old plants were examined under a bright-field microscope to capture images of stomata. The ImageJ software was utilised to quantify stomatal area. Three biological replicates from each line were used for all analyses. Error bars represent SE, with a two-way ANOVA showing p*<0.05, p**<0.01, p***<0.001, p****<0.0001.

### RPF2 interacts with RPL10A to regulate protein synthesis

To identify proteins that interact with AtRPF2, yeast two-hybrid (Y2H) screening was conducted using a cDNA library from Arabidopsis treated with mixed elicitors, leading to the identification of several potential AtRPF2-interacting proteins (Table S1). These included AtSOC1, AtNAC22, a Clavata complex interactor, and AtRPL10A. AtRPL10A, which plays a role in ribosome biogenesis and exhibits reduced nonhost disease resistance when silenced in *N. benthamiana* and Arabidopsis (Ramu et al., 2020), was chosen for further characterisation. The interaction between full length AtRPF2 and AtRPL10A was confirmed using a Y2H assay (Fig. 3A). *In silico* docking analysis indicated that the R210, R216, S190, F148, R7, K121, V213 amino acids of AtRPF2 may interact with F119, G120, G2, S104, E151, E217, R116 amino acids of AtRPL10A protein (Fig. 3B). Further the molecular dynamic simulations of both protein complexes confirm their transient interactions (Fig. 3C). *In planta* interactions of AtRPF2 and AtRPL10A was confirmed by bimolecular fluorescence complementation (BiFC) assay in *N. benthamiana* leaves by transient coexpression of *AtRPF2* fused to the N-terminal half of the enhanced green fluorescent protein (nGFP) and *AtRPL10A* fused to the C-terminal half of GFP (cGFP). Physical interaction of AtRPF2 and AtRPL10A reconstituted the expression of GFP in the nucleolus and nucleus (Fig. 3D). Additionally, the interactions were confirmed by biophysical methods using biolayer interferometry analysis with a His-tag sensor. The data showed that the *E. coli*-expressed AtRPF2-His protein associates with AtRPL10A-GST protein (Fig. 3E). *AtRPF2*-OE plants exhibited higher levels of 5.8S, 18S, and 25S rRNA than RNAi and Col-0 plants (Fig. 3F), with comparable RNA Integrity Numbers (RIN) (Fig. S5A-C). The ribosome profile of *AtRPF2*-OE and *AtRPL10A*-OE plants showed higher levels of 60S, 80S and polysomes when compared to Col-0 and RNAi lines (Fig. 3G&H). The *AtRPF2*-RNAi and *AtRPL10*A-RNAi plants also showed slightly higher ribosome profiles than Col-0, which may be due to redundant functions of other proteins (Fig. 3G&H). The translation capacity of these plants was evaluated using a radiolabeled 35S-[Met]-incorporation assay over 4 h in both Arabidopsis and *N. benthamiana*. *AtRPF2* overexpression plants showed increased protein synthesis compared to Col-0, whereas RNAi and mutant plants displayed significantly reduced protein synthesis levels (< 2-fold) compared to Col-0 plants (Fig. 3I & J). It is clear that, although RNAi plants have a higher ribosome profile than Col-0, translation is significantly affected. These findings indicate that *RPF2* downregulation impairs rRNA processing and affects protein synthesis.

**Figure 3.**
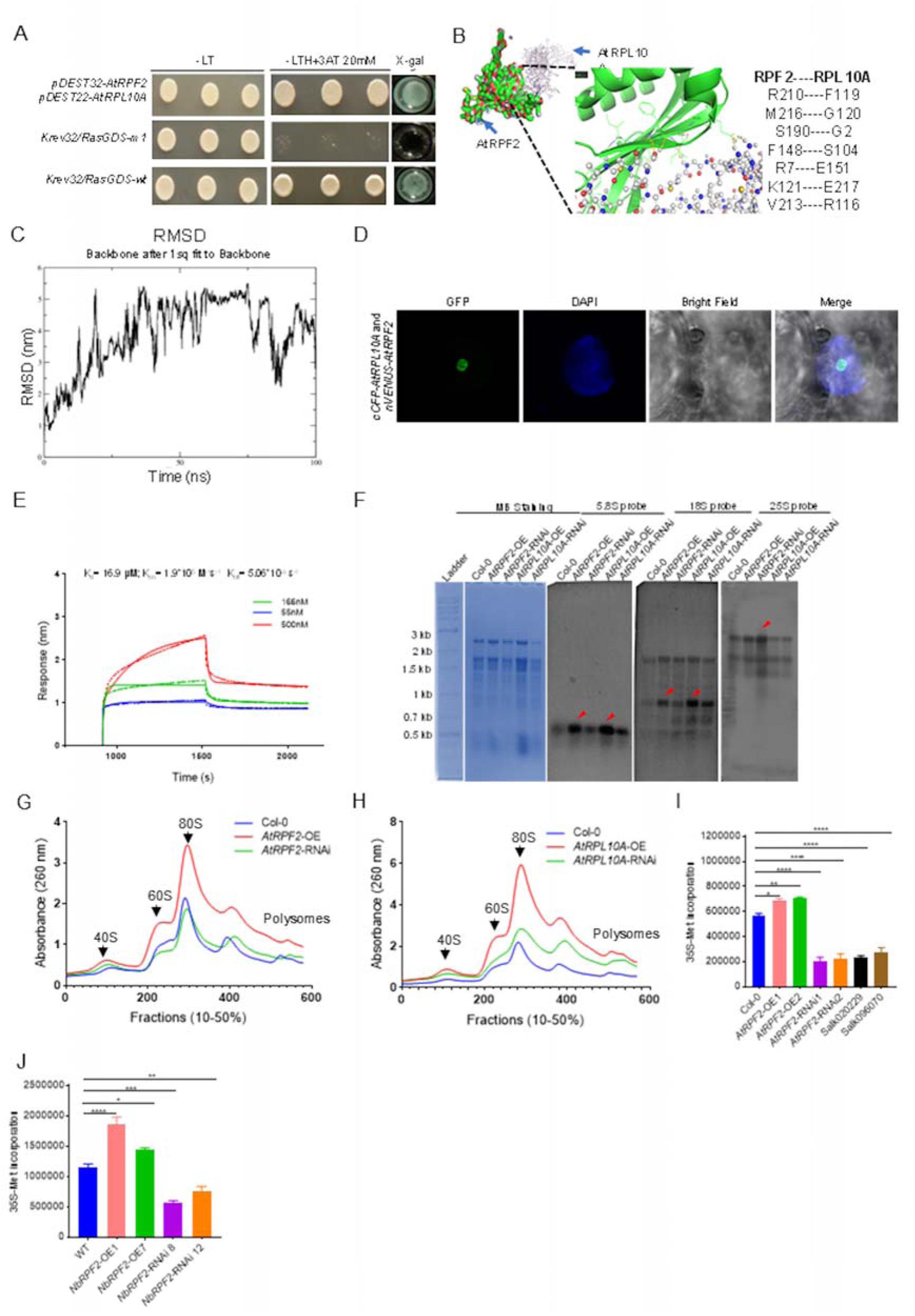
AtRPF2 associates with AtRPL10A to influence rRNA processing and protein production. A. Yeast-two-hybrid assay showing interaction of AtRPF2 and AtRPL10A. *AtRPL10A* in *pDEST32* and *AtRPF2* in *pDEST22* were expressed and co-transformed in *Mav203* yeast cells, then plated on double (-leu, -trp) and triple (-leu, -trp, -his with 20mM 3-AT) dropout media. *Krev1-RalGDS* and *Krev1-RalGDS-m1* used as positive and negative controls, respectively. X-Gal staining was performed to confirm the interactions. B. *In-silico* protein-protein docking demonstrated the interaction of AtRPF2 with AtRPL10A at different amino acids. The Alpha-fold modelled protein structures were used for docking studies using ClusPro 2.0. C. Molecular dynamic (MD) simulations indicated a stable interaction between AtRPL10A with AtRPF2 protein. The simulations were carried out using full-length proteins on the GrowMacs platform. D. *In planta* bi-molecular fluorescence complementation (BiFC) assay confirmed the interaction of AtRPF2 with AtRPL10A within the nucleus. The *pMDC-cCFP GFP-RPF2* or *pMDC-nVENUS GFP-RPL10A* constructs in Agrobacterium *GV3101* strain were co-expressed transiently in *N. benthamiana,* and after 48 h, the fluorescence was observed in confocal microscopy. DAPI was used to stain the nucleus. E. Bio-layer interferometry (BLI) assay verified the physical interactions of AtRPF2 with AtRPL10A. The *AtRPF2-*His in *pDEST15* and *AtRPL10A-*GST in *pDEST17* were expressed in *E. coli* and purified with their respective tags, and the His-tag sensor was used for BLI analysis. F. A Northern blot analysis revealed the accumulation of 5.8S, 18S, and 25S rRNA in *AtRPF2*-OE, *AtRPL10A*-OE, and RNAi Arabidopsis plants. Total RNA was extracted from four-week-old Arabidopsis leaves, immobilized on a nylon membrane, and probed with 5.8S, 18S, and 25S rRNA probes. G. Ribosome profiling from Col-0, *AtRPF2*-OE and *AtRPF2*-RNAi plants, H. Ribosome profiling from Col-0, *AtRPL10A*-OE and *AtRPL10A*-RNAi plants. Ten grams of tissues were used to extract ribosomes and 50-10% sucrose density gradient was used to profile. I. A 35S-methionine [35s-M] incorporation assay in Arabidopsis and J. in *N. benthamiana* plants demonstrating protein synthesis efficiency in overexpression and RNAi lines. A pulse chase assay with 35s-met incorporation into four-week-old leaf discs for 4 h was conducted. Methionine incorporation was quantified using a liquid scintillation counter. Error bars represent SE, with a two-way ANOVA showing p*<0.05, p**<0.01, p***<0.001, p****<0.0001.

### AtRPF2 and AtRPL10A regulate a unique set of protein synthesis

The RPF2 and RPL10A play roles in ribosome biogenesis and translation regulation. To identify differentially regulated proteins in Arabidopsis *AtRPF2*-OE, *AtRPL10A*-OE, *AtRPF2-*RNAi and *AtRPL10A*RNAi lines, comparative proteomic LC-MS/MS analysis was conducted on 4-week-old leaf tissues, using Col-0 as a control. Principal component analysis distinctly grouped the samples, showing no overlap when clustered using PC1 and PC2, with variations of 24.5% and 14.6%, respectively (Fig. 4A). The heat map clusters based on protein expression revealed that both *AtRPL10A*-OE and *AtRPF2*-OE plants, which were in the same clade, exhibited similar protein expression patterns, a trend also observed in both RNAi lines clustered in the same clade (Fig. 4B, & Fig. S6A). In *AtRPF2*-OE plants, 66 proteins were significantly upregulated, and 61 were downregulated compared to Col-0 plants (Fig. S6B, Supplementary Data S1, Sheets 1 and 2). In *AtRPF2*-RNAi plants, 60 proteins showed higher accumulation and 46 showed lower accumulation than in Col-0 (Fig. S6C, Supplementary Data S1, Sheet 3&4). Between the OE and RNAi lines, commonly 23 proteins were elevated, and 16 were downregulated (Supplementary Data S2, Sheets 1 and 2). Similarly, in *AtRPL10A*-OE plants, 23 proteins were upregulated and 38 were downregulated compared to Col-0 (Fig. S6D, Supplementary Data S3, Sheets 1 and 2) and in *AtRPL10A* RNAi lines, 33 proteins were upregulated and 47 downregulated compared to Col-0 (Fig. S6E, Supplementary Data S3, Sheet 3&4). Among these, five proteins were commonly upregulated in both *AtRPL10A*-OE and RNAi lines, and 10 were commonly downregulated compared to Col-0 (Fig. 4C, Supplementary Data S2, Sheet 3&4). Several proteins were also commonly differentially regulated in both *AtRPF2*-OE and *AtRPL10A*-OE lines compared with Col-0; however, the fold changes are higher in RPF2-OE plants (Table 1). Gene set enrichment analysis of upregulated proteins in *AtRPF2*-OE plants indicated an accumulation of proteins associated with the biosynthesis of secondary metabolites, amino acids, carbon, amino, and nucleotide sugars, the pyruvate metabolic pathway, and RNA degradation (Fig. 4D, Supplementary Data S4, Sheet 1). Increased levels of mitochondrial Cytochrome b-c1-comlex of subunit 10, caffeoyl-C6A-6-methyl-transferase, and UDP-glucuronic acid decarboxylase were more prevalent in *RPF2*-OE plants, resulting in elevated levels of glucosinolate metabolites. The glucosinolate pathway, tryptophan, and photosynthetic proteins were enriched, contributing to a strong phenotype with increased trichome density. Proteins involved in ATP-dependent protein folding, chaperoning, and NAD-binding activity (Fig. S6F) in the cytosol, mitochondria, plastids, and cell walls (Fig. S6G) were linked to plant water relations, ABA, carboxylic acid metabolism, and cold response, accumulating in *RPF2*-OE plants, potentially enhancing the superior phenotype (Fig. 4E). Conversely, in *RPF2*-RNAi lines, proteins associated with metabolism, amino acid biosynthesis, porphyrin metabolism, cofactors, and ribosome-related pathways were enriched among the downregulated proteins (Fig. S6H, Supplementary Data S4, Sheet 2). In RNAi lines, there was a reduction in RuBisCO accumulation factor 1.2, protoporphyrinogen oxidase 1, Multiple Organellar RNA Editing Factor 9 (MORF9) and 2, several ribosomal proteins, cytochrome P450 enzymes, translation initiation factor 2, and elongation factor G, which might explain the dwarf phenotype, decreased protein synthesis, and reduced glucosinolates in RNAi plants. In *AtRPL10A*-OE plants, proteins related to metabolic processes, secondary metabolite biosynthesis, glyoxylate, dicarboxylate, and carbon metabolism were enriched (Fig. 4F, Supplementary Data S4, Sheet 3). Photosynthesis-associated proteins (Fig. S6I) in the chloroplasts, cytosol, and mitochondria (Fig. S6J), involved in carboxylic acid metabolism, tryptophan, citrate, glyoxylate, and other organic acid metabolism, as well as water relations response (Fig. 4G), which contributed to the improved phenotype of *AtRPL10A*-OE plants. In *AtRPL10A*-RNAi lines, many proteins linked to metabolic, ribosomal, and secondary metabolite synthesis pathways were enriched (Fig. S6K, Supplementary Data S4, Sheet 4). Several ribosomal proteins, MORF9, cytochrome oxidases, 26S proteasome complex acetyl-CoA, Ferradoxin, Sulfur-E1 (Suf-E), peptidyl-prolyl cis-trans isomerases, peptidyl prolyl cis-trans isomerase FKBP15-2, and RNA-binding proteins were less accumulated than in Col-0, potentially affecting their growth and development. Proteomic analysis indicated that overexpression of *AtRPF2* or *AtRPL10A* resulted in increased protein levels involved in physiological processes, particularly carbon fixation, metabolism, and photosynthesis. In contrast, *AtRPL10A*-RNAi and *AtRPF2*-RNAi lines exhibited reduced proteins, which affected protein synthesis, GA levels, plant height, photosynthesis, and biomass.

**Figure 4.**
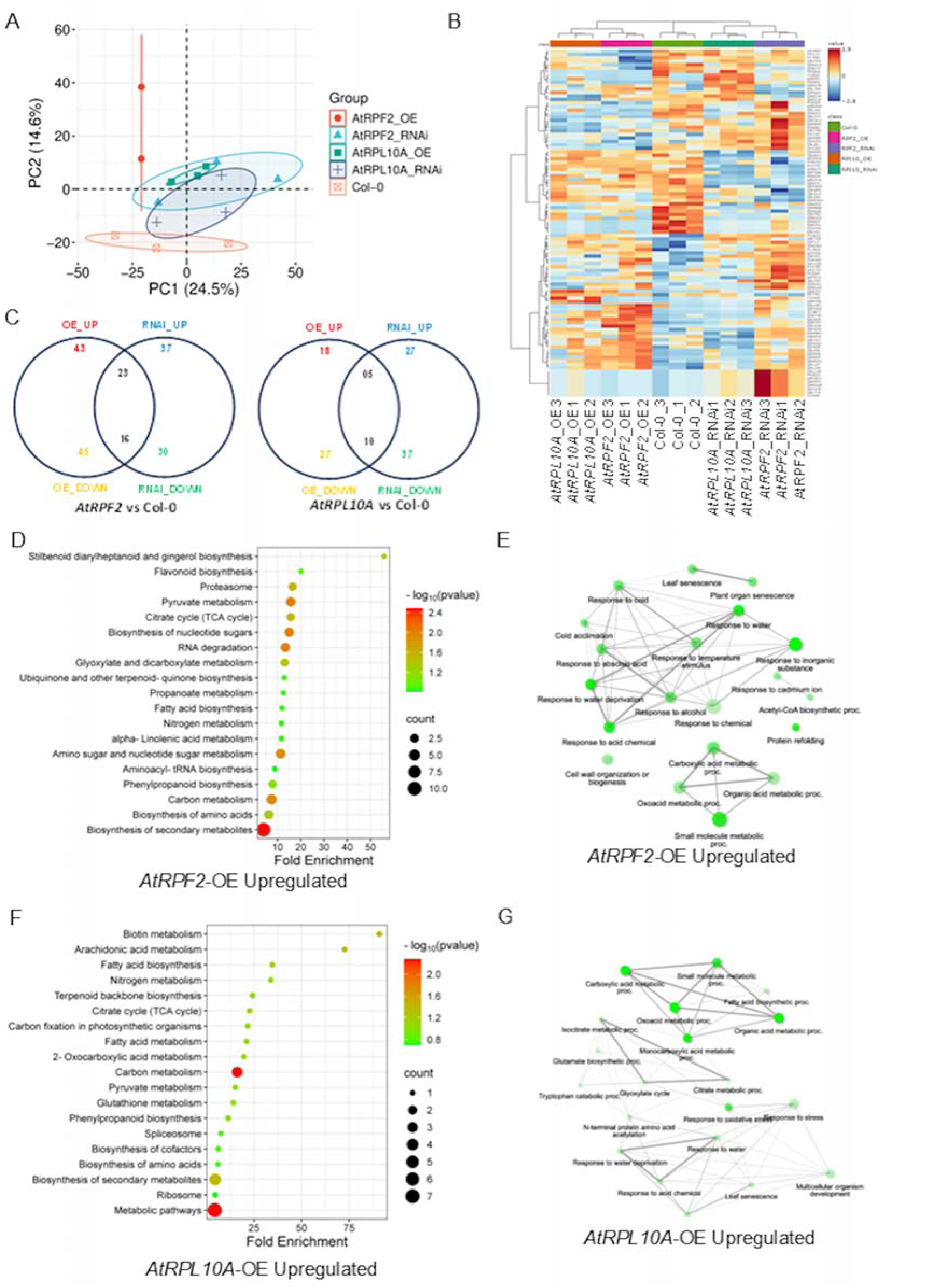
AtRPF2 and AtRPL10A are involved in regulation of protein synthesis. A. Proteomics data from *AtRPF2*-OE, *AtRPF2*-RNAi, *AtRPL10A*-OE, and *AtRPL10A*-RNAi lines under normal conditions. Total proteins from 4-week-old Arabidopsis plants’ leaf samples under normal conditions were developed using LC/MS- MS analysis. A) The differentially expressed proteins from each plant types are divided into PC1 and PC2 in a PCA plot, with the percentage for each PC indicated. B. Heat map illustrating the differentially expressed proteins from three biological replicates from OE and RNAi lines compared to Col-0. Clustering of the top 100 differentially expressed proteins was performed in SRplot analysis. C. Venn diagram depicting common and unique proteins from up and downregulated proteins of OE and RNAi lines compared to Col-0. Both up and down regulated proteins from each group were represented using the Venny 2.1 program. D. Gene set enrichment analysis of proteins upregulated in *AtRPF2*-OE plants showing different KEGG pathways. E. Network analysis of upregulated proteins demonstrating different biological functions in *AtRPF2*-OE plants. F. Gene set enrichment analysis of proteins upregulated in *AtRPL10A*-OE plants showing different KEGG pathways. G. Network analysis illustrating different biological functions of upregulated proteins in *AtRPL10A*-OE plants. The ShinyGO 0.82 network analysis tool was used to categorize the functions of proteins.

**Table 1.**
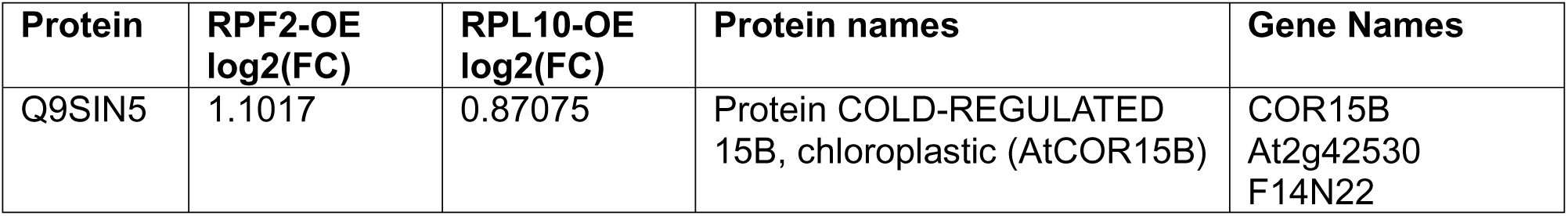
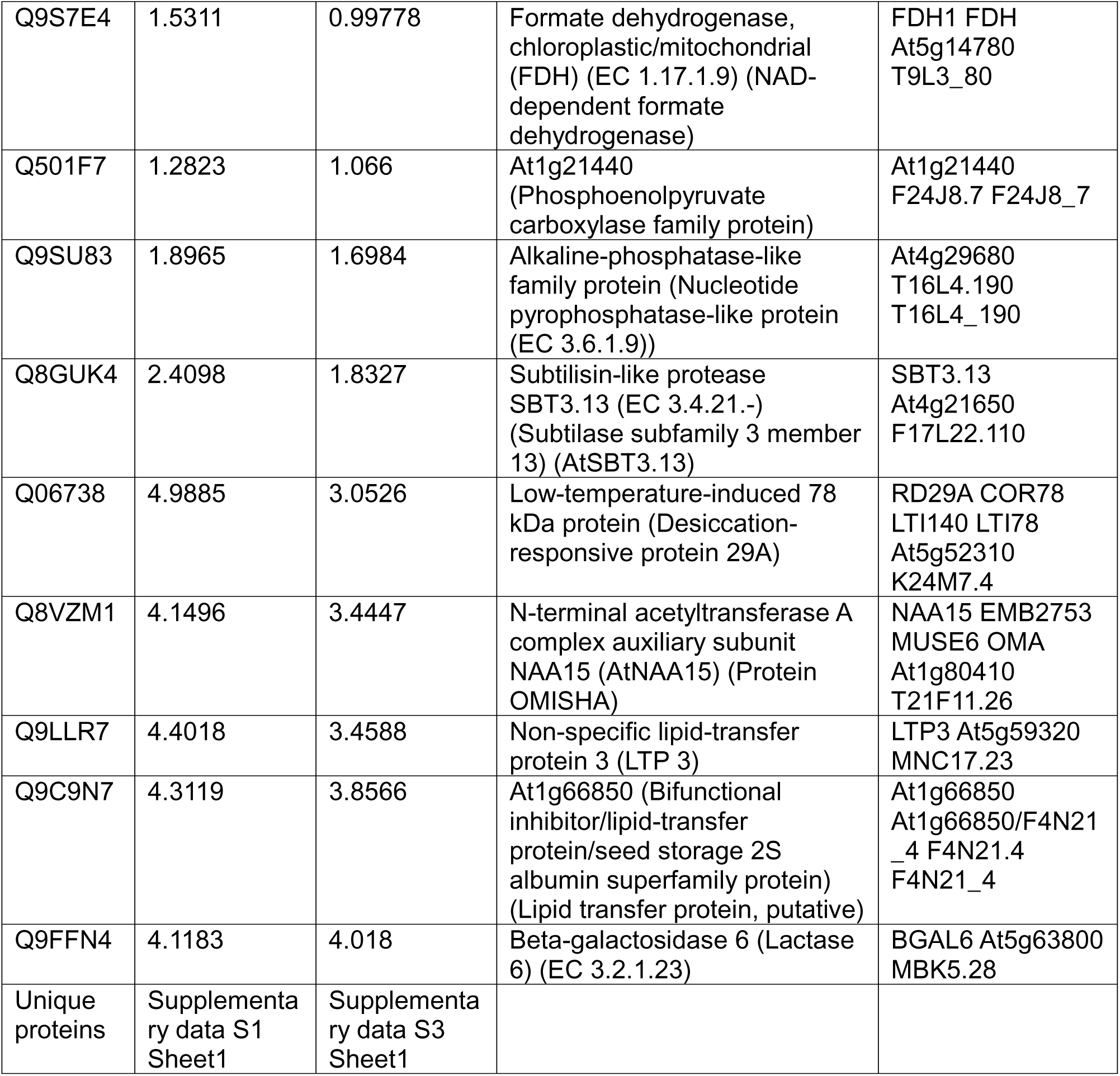
Common upregulated proteins in *AtRPL10A*-OE and *AtRPF2*-OE under control conditions.

| Protein | RPF2-OE<br>log<sub>2</sub>(FC) | RPL10-OE<br>log<sub>2</sub>(FC) | Protein names | Gene Names |
| --- | --- | --- | --- | --- |
| Q9SIN5 | 1.1017 | 0.87075 | Protein COLD-REGULATED<br>15B, chloroplastic (AtCOR15B) | COR15B<br>At2g42530<br>F14N22 |
| Q9S7E4 | 1.5311 | 0.99778 | Formate dehydrogenase, chloroplastic/mitochondrial (FDH) (EC 1.17.1.9) (NAD-dependent formate dehydrogenase) | FDH1 FDH<br>At5g14780<br>T9L3_80 |
| Q501F7 | 1.2823 | 1.066 | At1g21440 (Phosphoenolpyruvate carboxylase family protein) | At1g21440<br>F24J8.7 F24J8_7 |
| Q9SU83 | 1.8965 | 1.6984 | Alkaline-phosphatase-like family protein (Nucleotide pyrophosphatase-like protein (EC 3.6.1.9)) | At4g29680<br>T16L4.190<br>T16L4_190 |
| Q8GUK4 | 2.4098 | 1.8327 | Subtilisin-like protease SBT3.13 (EC 3.4.21.-) (Subtilase subfamily 3 member 13) (AtSBT3.13) | SBT3.13<br>At4g21650<br>F17L22.110 |
| Q06738 | 4.9885 | 3.0526 | Low-temperature-induced 78 kDa protein (Desiccation-responsive protein 29A) | RD29A COR78<br>LTI140 LTI78<br>At5g52310<br>K24M7.4 |
| Q8VZM1 | 4.1496 | 3.4447 | N-terminal acetyltransferase A complex auxiliary subunit NAA15 (AtNAA15) (Protein OMISHA) | NAA15 EMB2753<br>MUSE6 OMA<br>At1g80410<br>T21F11.26 |
| Q9LLR7 | 4.4018 | 3.4588 | Non-specific lipid-transfer protein 3 (LTP 3) | LTP3 At5g59320<br>MNC17.23 |
| Q9C9N7 | 4.3119 | 3.8566 | At1g66850 (Bifunctional inhibitor/lipid-transfer protein/seed storage 2S albumin superfamily protein) (Lipid transfer protein, putative) | At1g66850<br>At1g66850/F4N21_4 F4N21.4<br>F4N21_4 |
| Q9FFN4 | 4.1183 | 4.018 | Beta-galactosidase 6 (Lactase 6) (EC 3.2.1.23) | BGAL6 At5g63800<br>MBK5.28 |
| Unique proteins | Supplementary data S1 Sheet1 | Supplementary data S3 Sheet1 |  |  |

### AtRPF2 and AtRPL10A regulate plant defence proteins and improve resistance to host and nonhost pathogens

To investigate the molecular basis of pathogen defense responses in *AtRPF2, NbRPF2-*OE, and RNAi lines of *N. benthamiana*, these plants were syringe-infiltrated with the host pathogen *P. syringae* pv*. tabaci* and the nonhost pathogen *P. syringae* pv*. tomato* T1 at a concentration of 1 x 10^4^ cfu mL^-1^. The overexpression lines demonstrated enhanced resistance, whereas the RNAi lines were more susceptible to both host and nonhost pathogens (Fig. 5A-C). Pathogen proliferation in *NbRPF2*-RNAi lines was notably higher than that in wild-type plants, whereas *NbRPF2*-OE plants exhibited reduced pathogen growth (Fig. 5B&C). Additionally, Arabidopsis *AtRPF2*-OE and RNAi lines were exposed to the nonhost pathogen *P. syringae* pv*. tabaci*, and the host pathogen *P. syringae* pv*. tomato* (DC3000) through flood inoculation at 1 x 10^5^ cfu mL^-1^ concentrations. *AtRPF2*-OE plants displayed less growth of *P. syringae* pv*. tomato* (DC3000). In contrast, RNAi and mutant lines were susceptible to the host pathogen *P. syringae* pv*. tomato* (DC3000) (Fig. 5D). *ARPF2*-RNAi and mutant lines were also susceptible to the nonhost pathogen *P. syringae* pv*. tomato* T1 (Fig. 5E). Defense response genes were more upregulated in *AtRPF2*-OE plants than in Col-0 and RNAi lines. Specifically, *PDF1.2* showed a > 4-6 fold increase (Fig. 5F), *MYC1* > 4-5 fold (Fig. 5G), *ACD1* > 2-3 fold (Fig. 5H), and *PR5* > 23 fold upregulation (Fig. 5I) in *RPF2*-OE plants infected with the host pathogen compared with Col-0. In RNAi and mutant lines, these genes were either less induced or showed induction levels similar to those of Col-0 in response to the pathogen.

**Figure 5.**
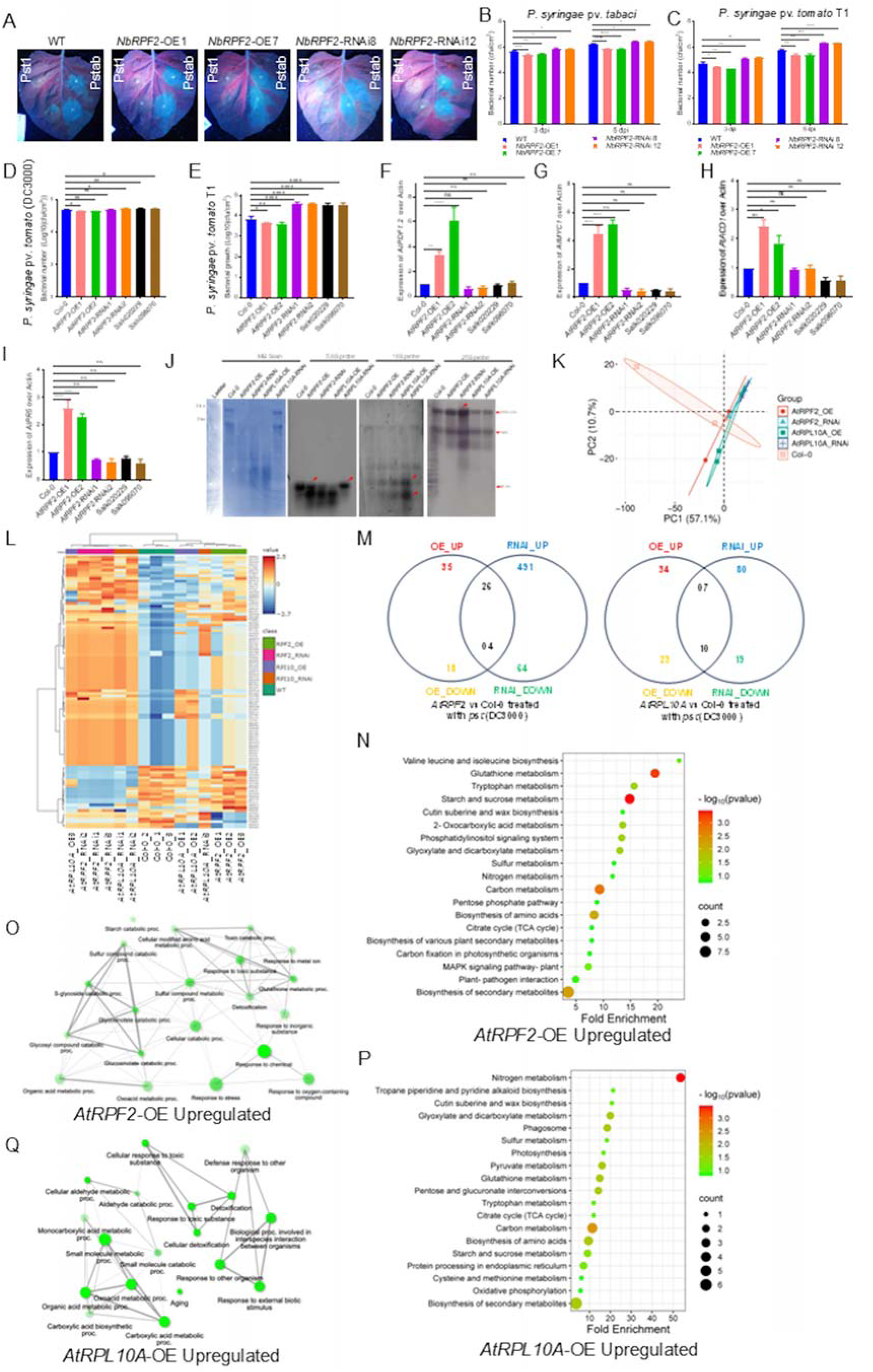
Response of RPF2-OE, RPF2-RNAi, RPL10A-OE, and RPL10A-RNAi plants to pathogen exposure. A. Observation of bacterial growth in host and nonhost species within *NbRPF2*-OE and *NbRPF2*-RNAi lines of *N. benthamiana* plants. Three weeks-old *N. benthamiana* plants were infiltrated using a needleless syringe with host *P. syringae* pv. *tabaci (Pstab)* and nonhost *P. syringae* pv. *tomato T1 (Pst1)* at a concentration of 1x10^7^ cfu/ml. B & C. Rate of bacterial growth for host and nonhost pathogens at 3-and 5-days post-inoculation (dpi). Leaves were inoculated with host and nonhost pathogens using a syringe. Average values from three biological replicates across three individual plants were calculated, and experiments were conducted three times with consistent results. Response of Arabidopsis *AtRPF2*-OE, *AtRPF2*-RNAi, and mutant plants to host and nonhost pathogens, including bacterial growth rates of D. *P. syringae* pv. *tomato* (DC3000), and E. *P. syringae* pv. *tomato* T1. The abaxial side of leaves was inoculated with host and nonhost pathogens (1 × 10^8^ cfu/ml) using a needleless syringe, and bacterial growth rates were assessed at 3 and 5 dpi. Gene expression analysis in response to host pathogen showing transcript levels of F. *PDF1*, G. *MYC2*, H. *ACD1*, I. *PR5* after 3 dpi. J. Northern blots showing *rRNA* profiling of *AtRPF2*-OE, *AtRPL10A*-OE, and RNAi lines in host pathogen-infected plants after 3 dpi. Total RNA from leaves infected with the host pathogen at 3 dpi was isolated and immobilized on a nylon membrane, and different 5.8S, 18S and 25S-rRNA probes were used. K. Proteomics data of *AtRPF2*-OE, *AtRPL10A*-OE, and RNAi lines in host-pathogen *P. syringae* pv. *tomato* (DC3000) infected Arabidopsis plants after 3 dpi are displayed in PC1 and PC2 on the PCA plot. L. Heat map illustrating the clustering of the top 100 differentially expressed proteins in pathogen-infected Arabidopsis plants. M. Venn diagram showing common and unique proteins from *AtRPF2*-OE, *AtRPL10A*-OE, and respective RNAi lines under pathogen infection conditions. N. Mapping of upregulated proteins from *AtRPF2*-OE in various KEGG pathways. O. Biological networks upon pathogen infection. P. Mapping of upregulated proteins from *AtRPL10A*-OE in various KEGG pathways. Q. biological networks after pathogen infection. The Science and Research online plot (SRplot) and ShinyGo 0.82 network analysis tools were utilized for mapping proteins in different categories.

Previously, we showed that plants overexpressing *AtRPL10A* exhibit resistance to host and nonhost pathogens compared with Col-0, whereas *AtRPL10A*-RNAi plants exhibit reduced resistance (Ramu et al., 2020). To examine the rRNA profile and protein translation in pathogen-infected *AtRPF2-*OE, *AtRPL10A*-OE, RNAi, and Col-0 plants, 4-week-old plants were flood-inoculated with *P. syringae* pv*. tomato* (DC3000). Analysis at 3 dpi showed that *AtRPL10A*-OE and *AtRPF2*-OE plants had increased 5.8S, 18S, and 25S rRNAs compared to Col-0, while RNAi lines showed decreased levels (Fig. 5J). Proteomic analysis identified differentially expressed proteins, with samples grouped using principal component analysis. Samples overlapped when clustered using principal components (PC1 and PC2), with variances of 57.1% and 10.7%, respectively (Fig. 5K). The heat maps showed that *AtRPF2*-OE and *AtRPL10A*-OE plants clustered in the same clade, whereas RNAi lines formed a separate clade (Fig. 5L & S7A). In *AtRPF2*-OE plants, 61 proteins were upregulated and 22 were downregulated (Fig. S7B, Supplementary Data S5, Sheet 1&2). In *AtRPF2*-RNAi lines, 517 proteins were upregulated and 68 were downregulated (Fig. S7C, Supplementary Data S5, Sheet 3&4). In *AtRPL10A*-OE plants, 41 proteins were upregulated, and 33 were downregulated (Fig. S7D, Supplementary Data S6, Sheet1&2). In *AtRPL10A*-RNAi lines, 87 proteins were upregulated and 29 were downregulated (Fig. S7E, Supplementary Data S6, Sheet 3 and 4). Many proteins were commonly regulated in both *AtRPF2/AtRPL10A* OE and RNAi lines (Fig. 5M, Table 2, Supplementary Data S7). Gene set enrichment analysis of *AtRPF2*-OE upregulated proteins showed involvement in secondary metabolites, carbon, starch, sucrose, glutathione metabolism, and amino acid biosynthesis (Fig. 5N). Key proteins, including Formate dehydrogenase (FDH), Superoxide dismutase (SOD), Glutathione S-transferase-F6, RNA-binding protein 8, and others, were upregulated in pathogen-treated *AtRPF2*-OE plants (Supplementary Data S8, Sheet 1). Proteins with various activities in molecular functions and cellular compartments (Fig. S7F, S7G), which are linked to glutathione, sulfur, glucosinolate, toxin catabolism, and stress response (Fig. 5O), potentially enhance the defense responses in *AtRPF2*-OE plants.

**Table 2.** Common upregulated proteins in *AtRPL10A*-OE and *AtRPF2*-OE under pathogen conditions.

| Protein | RPF2-OE<br>$\log_2(\text{FC})$ | RPL10-OE<br>$\log_2(\text{FC})$ | Protein names | Gene Names |
| --- | --- | --- | --- | --- |
| O81235 | 0.9393 | 1.2215 | Mn Superoxide dismutase [Mn] 1, mitochondrial (EC 1.15.1.1) (AtMSD1) (Protein MATERNAL EFFECT EMBRYO ARREST 33) | MSD1 MEE3<br>SODA<br>At3g10920<br>F9F8.26 |
| Q9SIB9 | 1.1395 | 1.321 | Aconitate hydratase 3, mitochondrial (Aconitase 3) (mACO1) (EC 4.2.1.3) (Citrate hydro-lyase 3) | ACO3<br>At2g05710<br>T3P4.5 |
| Q8LAS8 | 1.2987 | 1.7006 | S-formylglutathione hydrolase | SFGH |
|  |  |  | (AtSFGH) (EC 3.1.2.12)<br>(Esterase D) | At2g41530<br>T32G6.5 |
| P42645 | 1.5301 | 2.0834 | 14-3-3-like protein GF14 upsilon<br>(General regulatory factor 5) | GRF5<br>At5g16050 |
| Q96300 | 1.7775 | 2.1108 | 14-3-3-like protein GF14 nu<br>(General regulatory factor 7) | GRF7<br>At3g02520<br>F16B3.15 |
| P25853 | 1.9743 | 3.6463 | Beta-amylase 5 (AtBeta-Amy)<br>(EC 3.2.1.2) (1,4-alpha-D-glucan<br>maltohydrolase) (Protein<br>REDUCED BETA AMYLASE 1) | BAM5 BMY1<br>RAM1<br>At4g15210<br>FCAALL.97 |
| Q9FGS<br>0 | 2.8451 | 3.4886 | RNA-binding protein CP31B,<br>chloroplastic | CP31B<br>At5g50250 |
| P42770 | 3.132 | 2.7104 | Glutathione reductase,<br>chloroplastic (GR) (GRase) (EC<br>1.8.1.7) (Protein EMBRYO<br>DEFECTIVE 2360) | EMB2360<br>At3g54660<br>T5N23_20 |
| Q38946 | 3.3212 | 4.4962 | Glutamate dehydrogenase 2<br>(GDH 2) (EC 1.4.1.3) | GDH2<br>At5g07440 |
| Q8GUK<br>4 | 4.0128 | 3.6531 | Subtilisin-like protease SBT3.13<br>(EC 3.4.21.-) (Subtilase subfamily<br>3 member 13) (AtSBT3.13) | SBT3.13<br>At4g21650<br>F17L22.110 |
| O22788 | 4.5907 | 7.0452 | Probable peroxygenase 3<br>(AtPXG3) (EC 1.11.2.3)<br>(Caleosin-3) (Protein<br>RESPONSIVE TO<br>DESICCATION 20) | PXG3 CLO3<br>RD20<br>At2g33380<br>F4P9.15 |
| Unique<br>protein | Supplementar<br>y data S5<br>Sheet1 | Supplementar<br>y data S6<br>Sheet1 |  |  |

In the *AtRPF2*-RNAi lines, proteins were enriched in metabolic pathways, secondary metabolite biosynthesis, amino acid biosynthesis, ribosomes, and carbon metabolism (Fig. S7H). Several ribosomal proteins, such as eS25w, uS4c, uS5c, eL43z, uS13c, bS20c, bS18c, eS30z, bL21c, uS7cz, uL13y, eL6y, and eL37z, were downregulated. Additionally, chaperones, nitralases, pectin acetyltransferase, RuBisCO accumulation factor, dolichyl-diphosphooligosaccharide protein glycosyltransferase, protein DNA-damage inducible-1, and acyl carrier protein-3 were also downregulated, which contributed to increased disease susceptibility (Supplementary Data S8, Sheet 2). In *AtRPL10A*-OE plants, proteins were enriched in secondary metabolite biosynthesis, amino acid biosynthesis, carbon, nitrogen, pyruvate, and glutathione metabolism. Proteins related to glyoxylate, dicarboxylate, and phagosomes were enriched (Fig. 5P). In pathogen-treated *AtRPL10A*-OE plants, V-ATPase E1, SOD, Germin-like protein 3, 14-3-3, dehydrin cold-regulated 47 (COR47), Allen oxide cyclase 4, protease inhibitor, RNA-binding protein CP31P, Nudix hydrolase 3, Ran-binding protein homolog b, small coat protein complex II (COPII) GTPase SAR1B (Secretion Associated Ras Related GTPase 1B), Pectin esterase inhibitor 18, and senescence-associated protein B were upregulated (Supplementary Data S8, Sheet 3). Proteins enriched in plastids, mitochondria, and cell walls (Fig. S7I), along with glutamate dehydrogenase, protein disulfide reductase, metal binding, thioredoxin disulfide reductase, nitrilase, and antioxidant activity (Fig. S7J), are linked to defense responses, detoxification, ageing, and biotic stimuli (Fig. 5Q), thereby enhancing pathogen resistance. In *AtRPL10A*-RNAi cells, downregulated proteins were enriched in secondary metabolites, ribosomes, carbon metabolism, citrate cycle, glycolysis, and oxidative phosphorylation (Fig. S7K). Ribosomal proteins L6-2, uL6c, bS20c, S30, S25-4, and L9, as well as aspartate aminotransferase, spermidine synthase 2, glycolate oxidase 2, cef-G, GDP-mannose 3,5-epimerase, and V-ATPases were downregulated, contributing to disease susceptibility (Supplementary Data S8, Sheet 4). These findings indicate that RPF2 plays a distinct positive role alongside RPL10A in enhancing plant immunity against bacterial pathogens.

### H22AtRPF2 and AtRPL10A modulate drought tolerance through different mechanisms

To evaluate the response of *RPF2*-OE and RNAi lines to drought stress, *NbRPF2*-OE and *NbRPF2*-RNAi *N. benthamiana* plants were subjected to moisture stress at 50% field capacity (FC). Interestingly, *NbRPF2*-OE plants demonstrated enhanced drought resistance (Fig. 6A). These lines retained a stay-green appearance, whereas *NbRPF2*-RNAi and WT lines exhibited wilted leaves, indicating a drought-sensitive phenotype. During moisture stress, the loss of water through stomatal transpiration is a crucial factor linked to drought tolerance mechanisms. Detached leaves from *N. benthamiana NbRPF2*-OE plants maintained higher relative water content during drought stress (Fig. 6B). These findings imply that *RPF2* overexpression may increase ABA accumulation in plants, leading to reduced water loss and, thus, drought tolerance. RPL10A plays a role in ABA-dependent response as *Atrpl10a* mutants are insensitive to ABA (Ramos et al. 2020). The RPL10A was further examined by subjecting OE and RNAi plants to ABA-mediated seed germination and drought stress experiments. Germination of *AtRPF2*-OE seeds was delayed compared to Col-0 seeds when treated with ABA. The *AtRPF2*-RNAi and *Atrpf2* mutants were insensitive to ABA, germinating early (Fig. 6C), and a similar response was observed in *AtRPL0A*-RNAi plants (Ramos et al. 2020). The relative water loss in *AtRPF2*-OE plants was less than that in Col-0 and RNAi or mutant plants (Fig. 6D). Similarly, *AtRPL10*-OE plants had a reduced relative water loss than RNAi and Col-0 plants (Fig. 6E). Arabidopsis plants at 4 weeks old grown in pots were exposed to drought stress. *AtRPF2*-OE and *ARPL10A*-OE plants exhibited improved drought tolerance, whereas the RNAi lines were susceptible. The rRNA profile of *AtRPF2*-OE and *AtRPL10A*-OE plants under drought stress showed increased accumulation of 5.8S, 18S, and 25S rRNA compared to Col-0, whereas RNAi plants showed reduced levels of pre-rRNA accumulation (Fig. 6F).

**Figure 6.**
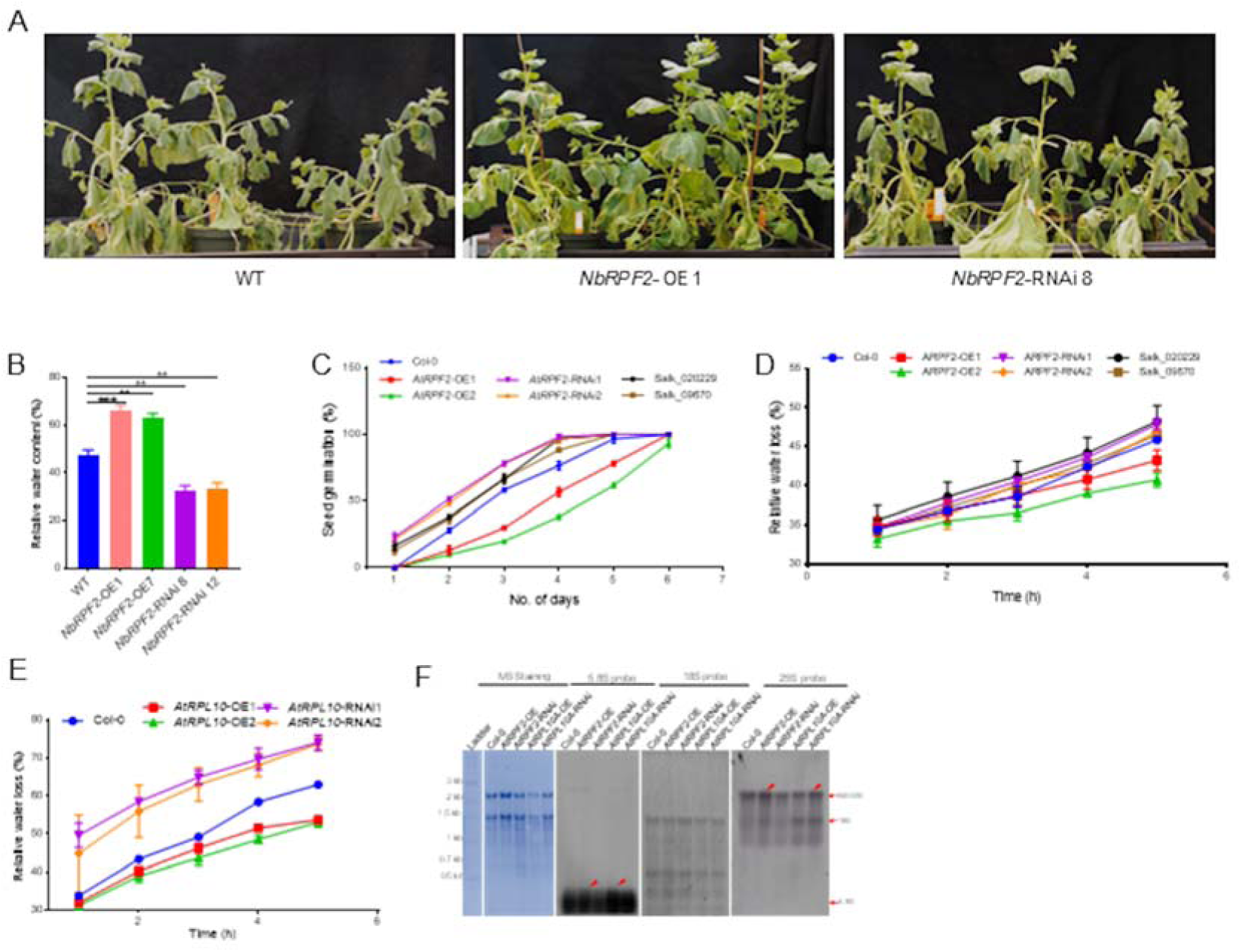
RPF2 and RPL10A play roles in ABA sensitivity and drought resistance. A. Six-week-old *N. benthamiana* plants exposed to drought stress showed wilting symptoms. *NbRPF2*-OE, *NbRPF2*-RNAi, and wild-type (WT) plants were exposed to progressive moisture stress at 50% field capacity (FC). The images were recorded after 4 days of plants reaching to 50% FC, B. The relative water content was evaluated in *NbRPF2*-OE and *NbRPF2-*RNAi lines in comparison to WT. C. ABA sensitivity assay showing delayed germination of *AtRPF2*-OE seeds and early germination of *AtRPF2*-RNAi. Ten seeds from each line were germinated on half-strength MS medium containing 1 µM ABA, and germination was monitored for 6 days. Relative water loss in D. *AtRPF2*-OE and *AtRPF2*-RNAi lines and E. *AtRPL10A*-OE and *AtRPL10A*-RNAi lines. The leaves from 4-week-old Arabidopsis plants were excised, air-dried, and weighed every hour. Three biological replicates from each line were analysed, and the experiments were repeated three times, yielding consistent results. Error bars represent SE, with a two-way ANOVA showing p*<0.05, p**<0.01, p***<0.001. F. Ribosomal RNA profiling from Arabidopsis plants under drought stress in *AtRPF2*-OE, *AtRPL10A*-OE, and RNAi lines. Four-week-old Arabidopsis plants were subjected to drought stress at 50% field capacity for four days, after which samples were collected, and total RNA was extracted and immobilised on a nylon membrane for probing with 5.8S, 18S, and 25S rRNA probes.

Comprehensive proteomic analysis was conducted on *AtRPF2*-OE, *AtRPL10A*-OE, and their respective RNAi lines under drought conditions, given the roles of RPF2 and RPL10A in ribosome assembly. Proteins with differential expression were categorized using principal component analysis and organized into clusters based on PC1 and PC2, which accounted for 20.9% and 13.3% of the variation, respectively (Fig. 7A). Heatmap analysis revealed that *AtRPF2*-OE and *AtRPL10A*-OE proteins formed distinct clusters, whereas the RNAi lines shared a cluster with *AtRPF2*-OE proteins under drought conditions (Fig. 7B and S8A), indicating separate tolerance mechanisms. In *AtRPF2*-OE plants, 89 proteins were upregulated and 135 were downregulated compared to Col-0 under drought stress (Fig. S8B, Supplementary Data S9, Sheets 1 and 2). *AtRPF2*-RNAi plants exhibited 103 upregulated and 170 downregulated proteins (Fig. S8C, Supplementary Data S9, Sheet 3&4). *AtRPL10A*-OE plants had 61 upregulated and 219 downregulated proteins (Fig. S8D, Supplementary Data S10, Sheet 1&2), whereas *AtRPL10A*-RNAi plants showed 41 upregulated and 109 downregulated proteins (Fig. S8E, Supplementary Data S10, Sheet 3&4). Some proteins were commonly regulated between the OE and RNAi lines of *AtRPF2* and *AtRPL10A* (Supplementary Data S11), with both unique and shared regulations across different lines (Fig. 7C). Although unique proteins accumulated under drought conditions, many proteins were regulated by both AtRPF2 and AtRPL10A (Table 3). In *AtRPF2*-OE plants, proteins associated with secondary metabolite biosynthesis, ribosomes, carbon metabolism, glucosinolate biosynthesis, photosynthesis, carotenoid, and oxidative phosphorylation were enriched (Fig. 7D). In plants with overexpressed genes, several ribosomal proteins from large and small subunits, eukaryotic initiation factor-4A (eIF4-A), mRNA binding proteins, and Receptor for Activated C Kinase 1 (RACK1) were increased. Proteins in Photosystem I and II, chlorophyll synthase, exokinase1, Plasma membrane Intrinsic Protein (PIP2-1), heat shock protein (hsp70), DnaJ Homolog Subfamily C Member 3, H+ ATPase, and Aldehyde dehydrogenase (ALDH3) were upregulated, enhancing drought tolerance (Supplementary Data S12, Sheet 1). Proteins with RNA-binding and molecule-binding capabilities (Fig. S8F) in the cell wall, cytosol, and polysomes (Fig. S8G), which are linked to translation, vesicle transport, photosynthesis, and stress response, were upregulated (Fig. 7F), thereby aiding drought resistance. In RNAi plants, downregulation of proteins in metabolic pathways, oxidative phosphorylation, glutathione, ribosomes, and MAPK signaling indicates increased drought susceptibility (Fig. S8H). Drought vulnerability results from the downregulation of feradoxin1, COR15-A, B, ERD10, ClpB3, glutathione reductase, eIF2-beta, and other proteins (Supplementary Data S12, Sheet 2). In *AtRPL10A*-OE plants, proteins associated with metabolic processes and carbon metabolism were enriched (Fig. 7E). Proteins, including dolichyl-diphospho oligosaccharide, glycosyltransferase, dnaJ, OMT1, eIF3-F, and others were upregulated, contributing to drought tolerance (Supplementary Data S12, Sheet 3). Proteins with nucleotide-binding activity (Fig. S8I) in the cytosol and ribosomes (Fig. S8J) are linked to water transport and the MAPK cascade (Fig. 7G), thereby enhancing translation under drought conditions. In *AtRPL10A*-RNAi plants, downregulated proteins were associated with ribosomes, RNA degradation, and carbon metabolism, reducing drought resistance (Fig. S8K). Downregulation of ERD10, ClpB3, V-H+-ATPase, ribosomal proteins, and other proteins compared to Col-0 (Supplementary Data S12, Sheet 4) contributed to drought susceptibility.

**Figure 7.**
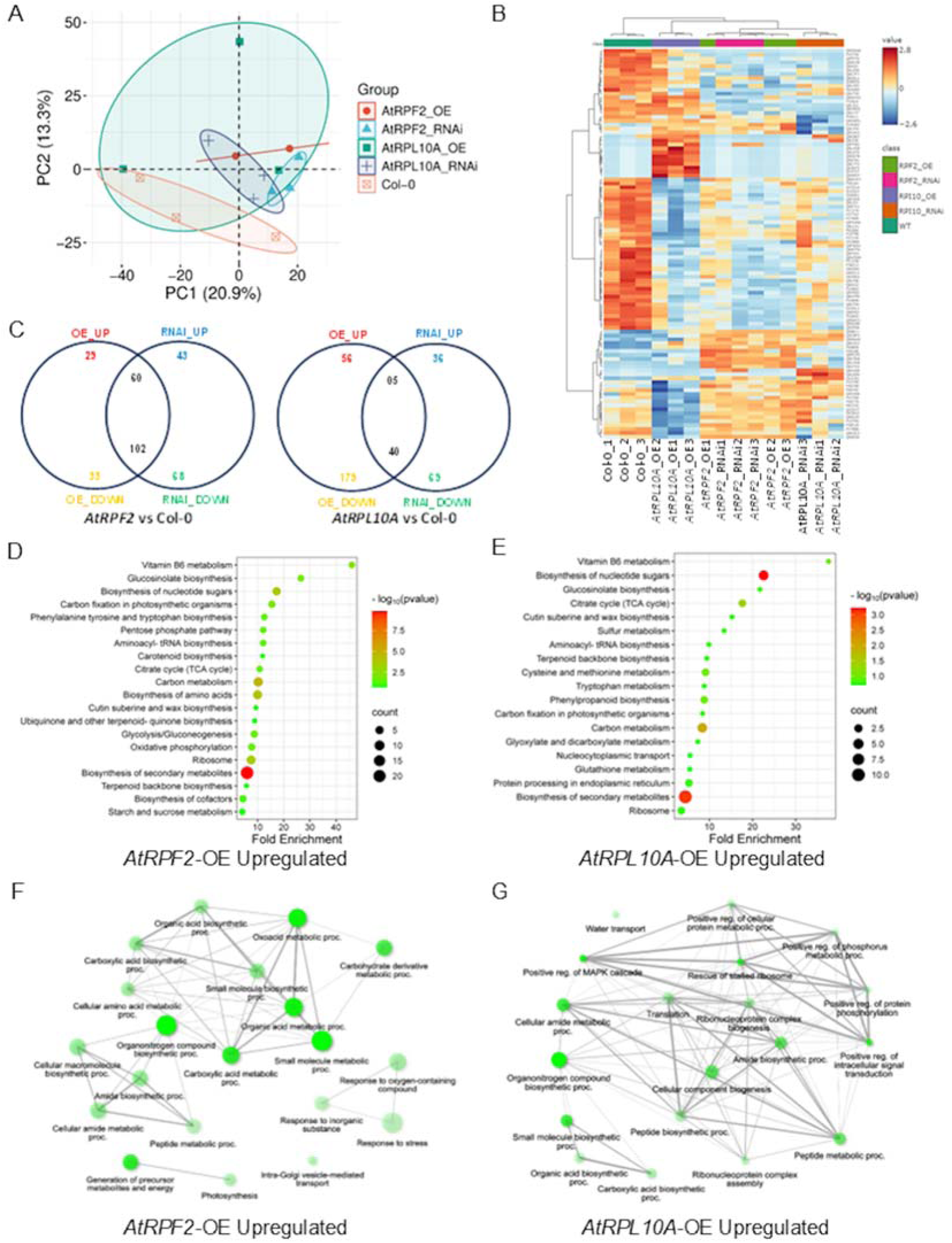
AtRPF2 and AtRPL10A have unique functions in regulating the translation of proteins that respond to drought stress in Arabidopsis. A. The PCA plot showing the proteomics data that differed in *AtRPF2*-OE, *AtRPL10A*-OE, and RNAi lines under drought conditions compared to Col-0, categorised in PC1 and PC2. B. Heatmap showing the differentially expressed proteins from each line clustering in different groups under drought stress. The top 100 differentially expressed proteins were mapped. C. The proteins that were either commonly or uniquely upregulated or downregulated in *AtRPF2-*OE and *AtRPL10A*-OE or RNAi plants over Col-0 plants are presented through Venn diagram. D. KEGG pathways and E. biological networks illustrating the proteins that are upregulated in *AtRPF2*-OE and *AtRPL10A*-OE plants, F and G. Molecular functions of upregulated proteins from *AtRPF2*-OE and *AtRPL10A*-OE plants. The SRplot analysis and ShinyGO 0.82 network analysis tools were used to classify these proteins.

**Table 3.** Common upregulated proteins in *AtRPL10A*-OE and *AtRPF2*-OE under drought condition.

| Proteins | RPF2-OE<br>log<sub>2</sub>(FC) | RPL10-OE<br>log<sub>2</sub>(FC) | Protein names | Gene Names |
| --- | --- | --- | --- | --- |
| Q944K2 | 0.94833 | 1.0279 | Dolichyl-diphosphooligosaccharide-<br>protein glycosyltransferase 48 kDa<br>subunit (Oligosaccharyl transferase<br>48 kDa subunit) (Protein<br>DEFECTIVE GLYCOSYLATION 1) | OST48 DGL1<br>At5g66680<br>MSN2.7 |
| F4J9G2 | 1.2086 | 0.89132 | Rhodanese/Cell cycle control<br>phosphatase superfamily protein | At3g59780 |
| Q94AW8 | 1.6921 | 1.232 | Chaperone protein dnaJ 3 (AtDjA3)<br>(AtJ3) | ATJ3 A3 J3<br>At3g44110 |
| Q9SGE0 | 1.7291 | 1.7508 | UDP-D-apiose/UDP-D-xylose<br>synthase 2 | AXS2<br>At1g08200 |
| Q9FIE8 | 1.8665 | 2.6558 | UDP-glucuronic acid<br>decarboxylase 3 (EC 4.1.1.35) | UXS3<br>At5g59290 |
|  |  |  | (UDP-XYL synthase 3) (UDP-glucuronate decarboxylase 3) (UGD) (UXS-3) | MNC17.20 |
| Q8VZM1 | 2.9401 | 3.3718 | N-terminal acetyltransferase A complex auxiliary subunit NAA15 (AtNAA15) (Protein OMISHA) | NAA15<br>EMB2753<br>MUSE6 OMA<br>At1g80410 |
| Q9C5N2 | 3.1467 | 3.1616 | Transmembrane 9 superfamily member 9 (Endomembrane protein 2) (AtTMN9) | TMN9 EMP2<br>At5g25100<br>T11H3_110 |
| Q9MAH0 | 4.2449 | 4.2931 | Phosphoenolpyruvate carboxylase 1 (AtPPC1) (PEPC 1) (PEPCase 1) (EC 4.1.1.31) (107-kDa PEPC polypeptide) | PPC1 p107<br>At1g53310<br>F12M16.21 |
| O04630 | 4.6685 | 3.1513 | Threonine--tRNA ligase, mitochondrial 1 (EC 6.1.1.3) (AtSYT1) (Threonyl-tRNA synthetase) (ThrRS) | THRRS<br>At5g26830<br>F2P16.7 |
| O22788 | 4.7765 | 2.8007 | Probable peroxygenase 3 (AtPXG3) (EC 1.11.2.3) (Caleosin-3) (Protein RESPONSIVE TO DESICCATION 20) | PXG3 CLO3<br>RD20<br>At2g33380 |
| P56808 | 6.1382 | 3.9545 | Small ribosomal subunit protein uS19c (30S ribosomal protein S19, chloroplastic) | rps19<br>AtCg00820 |
| Q9C522 | 6.3925 | 5.8911 | ATP-citrate synthase beta chain protein 1 (ATP-citrate synthase B-1) (EC 2.3.3.8) (ATP-citrate lyase B-1) (Citrate cleavage enzyme B-1) | ACLB-1<br>At3g06650<br>F5E6.2 T8E24.7 |
| Unique protein | Supplementary data S9 Sheet1 | Supplementary data S10 Sheet1 |  |  |

## Discussion

Plants activate complex defense mechanisms upon detecting pathogens, but many bacterial pathogens can suppress or manipulate these responses, leading to disease. While host resistance often depends on specific resistant (*R*) genes, nonhost resistance involves multiple genes and broader defense layers (Fonseca & Mysore, 2019; Gill et al., 2015; Senthil-kumar, 2013).

Through a VIGS-mediated forward genetics screen, we previously identified several genes that play roles in nonhost disease resistance (Kaundal et al., 2017; Lee et al., 2017; Rojas et al., 2012; Wang et al., 2012). Many ribosomal protein-encoding genes, such as *RPL12, RPL19, RPL10A, RPL10B, RPS30, RPL30,* and *RPL23,* have been linked to both host and nonhost resistance (Nagaraj et al., 2016; Ramu et al., 2020). Several ribosomal protein-encoding genes exhibit differential expression during *Xanthomonas* infection of rice (Pal et al., 2022; Solano et al., 2019; Vemanna et al., 2019). The role of translation mechanisms, particularly ribosome biogenesis and ribosomal proteins, in plant defense remains poorly understood. Ribosome biogenesis is a meticulously coordinated and involves the conversion of 45S rRNA into 18S, 5.8S, and 25S rRNAs, which are integral components of the large and small subunits required for mRNA translation.

Overexpressing *RPF2* in *N. benthamiana* and Arabidopsis resulted in enhanced growth, attributed to the rearrangement of leaf phyllotaxy, which improves light perception, photosynthesis, and biomass accumulation (Gommers et al., 2013; Vemanna et al., 2017). The role of *RPF2* in modifying the plant leaf morphology and 5.8S rRNA processing was reported (Choi et al., 2020; Maekawa et al., 2018). Another contributing factor could be the increased stomatal aperture, which enhances CO_2_ uptake, thereby facilitating photosynthesis and biomass production. Additionally, the higher trichome density observed in both Arabidopsis and *N. benthamiana* OE plants may play a crucial role in reducing water loss and bolstering plant defense against herbivores and pathogens (War et al., 2012). The elevated trichome levels in overexpression lines might be linked to the accumulation of gibberellic acid (GA) and salicylic acid (SA), with GA promoting trichome development (Zang et al., 2023; Perazza et al., 1998).

Although RPF2 is conserved in eukaryotes, plant RPF2 belongs to a distinct clade separate from fungi and animals, suggesting it may have functions beyond those observed in yeast. Notably, it interacts with plant-specific NAC and SOC transcription factors (Table S1). Further research is necessary to elucidate these interactions and their functional significance in plants. Here, we show that AtRPF2 associates with AtRPL10A, which plays a role in 60S ribosome biogenesis in the nucleus, suggesting their potential involvement in ribosome assembly. In yeast, RPF2 is known to interact with 5srRNA, RPL5, RPL11, and RRS1, all of which are adjacent to RPL10 in the ribosome complex. These components are crucial for processing pre-rRNA into 25S rRNA and for the maturation of the 60S subunit (Zhang et al., 2007; Maekawa et al., 2018). The interaction between AtRPF2 and AtRPL10A is vital for processing the pre-ribosome subcomplex, particularly 27S pre-rRNA, and for exporting the large subunit from the nucleus (Zhang et al., 2007). Overexpression of *AtRPF2* and *AtRPL10A* has demonstrated their involvement in rRNA processing and ribosome biogenesis, as evidenced by the accumulation of 25S, 18S, and 5.8S rRNA and a normal ribosome profile. Mutants exhibited reduced rRNA accumulation, potentially hindering rRNP assembly in ribosomes and affecting ribosome biogenesis, as indicated by the ribosome profile. AtRPL10A interacts with other ribosomal proteins, including L23, L30, S30, and RRM proteins (Ramu et al., 2020). Both RPF2 and RPL10A regulate protein synthesis, thereby influencing plant growth in Arabidopsis and *N. benthamiana*. *AtRPF2*-OE and *AtRPL10A*-OE plants display similar differentially expressed proteins in neighbouring clades; however, some proteins accumulate differently in each, suggesting their independent roles in mRNA translation regulation. The extra-ribosomal functions of AtRPL10A, through interactions with MYB transcription factors, regulate photosynthesis and defence response genes (Zorzato et al., 2015, and Ferreira et al., 2025). Similar mechanisms may be hypothesised for AtRPF2 due to its interactions with plant-specific NAC and SOC1 transcription factors. AtRPF2 influences chaperones, protein folding, lipid, and NAD-binding proteins involved in cell wall, ABA, water deprivation, and chemical responses across developmental stages, including those from plastids, chloroplasts, mitochondria, and transmembrane proteins. AtRPL10A regulates oxidoreductases, iron-sulfur cluster binding, acyl carriers, and antioxidant proteins involved in metabolic pathways, oxidative stress responses, and water deprivation-related proteins in mitochondria, chloroplasts, and the cytosol.

*AtRPL10A* regulates plant immunity through basal tolerance mechanisms at transcriptional and translational levels (Ramu et al., 2020). Silencing of *NbRPF2* in *N. benthamiana* and *Atrpf2* Arabidopsis mutants compromised resistance to host and nonhost pathogens. *NbRPL12* and *NbRPL19* silencing in *N. benthamiana* weakens disease resistance (Nagaraj et al., 2016). *AtRPF2*-RNAi and *AtRPL10A-*RNAi lines were susceptible to host pathogen *P. syringae* pv. *tomato* DC3000 and nonhost pathogen *P. syringae* pv. *tomato* T1, while overexpressing plants showed resistance to DC3000, indicating their role in basal and PAMP-triggered immunity (Fig 5 D&E; Ramu et al., 2020). Quantitative proteomics analysis showed unique proteins accumulating in *AtRPF2*-OE and *AtRPL10A*-OE plants, however, AtRPF2 regulates more proteins during infection. *AtRPL10A*-RNAi lines showed more downregulated genes (Ramu et al., 2020). MORFs, upregulated in both *AtRPL10*-OE and *AtRPF2*-OE lines, affect RNA maturation and organellar development (Ramu et al., 2020; Zhang et al., 2021). Disease tolerance and biomass increase may result from higher accumulation of photosynthesis proteins (Open et al., 2020; Zhang et al., 2024). In *RPF2*-RNAi plants, downregulation of translation factors and proteins correlated with dwarf phenotypes. Reduced secondary metabolites and lower glucosinolate accumulation in leaves and trichomes of *RNAi* plants affect plant defence mechanisms (Frerigmann et al., 2012; Lv et al., 2022). For example, secondary metabolites such as phytosterols are reduced in RNAi plants which are known to enhance plant innate immunity against bacterial pathogens (Wang et al., 2012).

In Arabidopsis plants overexpressing *AtRPL10A* or *AtRPF2*, proteins like SOD, aconitase, and 14-3-3 accumulate to stabilize H+ ATPase in plant defense. Stabilization of 14-3-3, with decreased *GCN4* expression during pathogen invasion, increases disease susceptibility (Kaundal et al., 2017). In *AtRPF2*-RNAi and *AtRPL10A*-RNAi plants, ribosomal proteins including es25w, us4c, uL5c, eL43z, uS13c, bS20c, bS18c, eS30z, bL21c, uS7cz, uL13y, eL6y, eL37z L6-2, ul6c, S25-4, and L9 are downregulated during pathogen exposure in *N. benthamiana* (Ramu et al., 2020). Similar downregulation occurs in *Fusarium oxysporus* sp*. vanillae* and rice infected with *X. oryzae* pv*. oryzae* (Pal et al., 2024; Solano de La cruz et al., 2019). In Arabidopsis *RPF2*-overexpression plants, upregulated peptidyl-prolyl cis-trans isomerases known to provide pathogen resistance (Mokryakova et al., 2014). The RNA recognition motif (RRM) protein, upregulated in *AtRPF2* overexpression plants, aids defense and interacts with AtRPL10A (Ramu et al., 2020). *AtRPF2* overexpression increases allene oxidase levels, involved in jasmonic acid synthesis and defense. Higher SA levels indicate activation of systemic acquired resistance (Park et al., 2007). In *AtRPL10A* overexpression plants, resistance occurs through SA-independent mechanisms (Ramu et al., 2020), due to specific mRNA translation by RPL10A proteins (Ferreyra et al., 2010; Carvalho et al., 2008; Zarzatto et al., 2015), suggesting both RPL10A and RPF2 operates through independent pathways in immunity.

Plants overexpressing *AtRPF2* were sensitive to ABA and exhibited drought tolerance through reduced water loss, despite larger stomatal openings (Galdon-armero et al., 2018; Mo et al., 2015). While *Atrpl10a* mutants don’t respond to ABA, *AtRPL10A*-OE plants showed drought tolerance (Ramos et al., 2020), similar to *AtRPF2*-OE lines. In *AtRPF2*-OE plants, upregulated RACK1 contributed to ABA and stress responses. The Arabidopsis *Atrack1* mutant may impair ribosome translation due to decreased water content (Guo et al., 2011). Ribosomal proteins regulate protein synthesis and plant stress responses (Dawane et al., 2024). *AtRPF2*-OE and *AtRPL10A*-OE plants showed upregulation of photosystem II, I, chlorophyllase, and other key proteins. Under drought, FNR, NPQ4, and RAF2 are maintained in drought-resistant rice plants (Dawane et al., 2024). Translation initiation genes are shown to enhance translation of metabolic genes (Weiss et al., 2021).

In *AtRPF2-OE* and *AtRPL10A-OE* plants, ROS-scavenging proteins like dehydrogenases, HSPs, ALDH, and AKR proteins were upregulated to combat oxidative stress. Previous research showed ALDH and AKRs scavenge ROS and reactive carbonyl compounds (Niranjan et al., 2021; Vemanna et al., 2017). *AtRPF2*-overexpressing plants accumulated more ribosomal proteins than *AtRPL10A*-overexpressing plants. RNAi plants showed downregulation of ribosomal proteins, eIFs, and other essential proteins, increasing drought susceptibility. *AtRPF2*-RNAi plants exhibited downregulation of dehydrins, stress-response proteins, and chaperonins. Dehydrins maintain protein function during stress (Babitha et al., 2013, 2015). Downregulated proteins in *AtRPL10A*-RNAi lines increased oxidative stress susceptibility, while SOD or HSP overexpression enhances stress response (Salehi et al., 2020; Vemanna et al., 2017). AtRPF2 is involved in rRNA processing and interacts with AtRPL10A to regulate ribosome biogenesis and the translation of specific mRNAs. AtRPF2 and AtRPL10A play an integral role in regulating plant growth processes and conferring resistance to biotic and abiotic stresses, making them promising candidates for the development of climate-resilient crops.

## Materials and methods

### Plant materials and bacterial strains

In this study, Arabidopsis Col-0 and *N. benthamiana* were used. The *Agrobacterium tumefaciens* strain GV3101, which expresses *TRV2::19A10* (*NbME19A10* http://vigs.noble.org), was cultivated at 28 °C in LB medium with rifampicin and kanamycin. The VIGS procedure involved using TRV1- and TRV2-expressing *Agrobacterium* strains in a 1:1 ratio, which were infiltrated into four-week-old *N. benthamiana* leaves using a needleless syringe (Senthilkumar & Mysore, 2014). Full-length *NbRPF2* from *N. benthamiana* and AtRPF2 or AtRPL10A from Arabidopsis were PCR-amplified from cDNA and cloned under the *CaMV*-35S promoter in the *pH7GWIWG2*(II) binary vector (Karimi et al., 2007). Similarly, constructs for *NbRPF2*-RNAi and *AtRPF2*-RNAi were created using the *pB7GWIWG2*(I) binary vector. Homozygous lines of Arabidopsis T-DNA insertion mutants (*Atrpf2)*, *salk-020229* and *salk-96070*, were obtained from the Arabidopsis Biological Resource Center (ABRC) for further study. Transgenic plants were developed using *Agrobacterium*-mediated transformation with *N. benthamiana* leaf discs as explants and the floral-dip method for Arabidopsis, and these were screened on hygromycin (25 μg/ml) and Basta (25 μm). Previously developed Arabidopsis *AtRPL10A* overexpression and RNAi lines were used (Ramu et al., 2020). *Pseudomonas syringae* pv*. tomato* (DC3000), *Pseudomonas syringae* pv*. tomato* T1, and *Pseudomonas syringae* pv*. tabaci* were grown in King’s B medium at 30 °C with the antibiotics Rifampicin (10µg/ml), Kanamycin (50µg/ml).

### Bacterial disease assays in *N. benthamiana* and Arabidopsis

To evaluate the infection assays for both the host pathogen (*P. syringae* pv*. tabaci*) and the nonhost pathogen (*P. syringae* pv*. tomato* T1), three-week-old *N. benthamiana* plants with silenced and overexpressed *NbRPF2* were vacuum-infiltrated with bacteria expressing *GFPuv* (Wang et al., 2007). Samples were collected from three biological replicates at 0-, 3-, and 7-days post-inoculation (dpi) to determine the bacterial titer on King’s B agar medium supplemented with suitable antibiotics. After two days of incubation at 28 °C, bacterial colonies were counted. To evaluate bacterial population host or nonhost pathogen was syringe-infiltrated into the leaves of *N. benthamiana*. Four-week-old Arabidopsis plants cultivated on Murashige and Skoog (MS) plates were flood-inoculated with host or nonhost pathogen for one minute using 40 ml of bacterial suspension as described (Ishiga et al., 2011). Bacterial symptoms were documented, and tissues were collected 3 and 5 days after infection to measure bacterial proliferation from the entire rosette leaves, as described previously (Ishiga et al., 2017). For each treatment, three biological replicates were collected from the leaves using a cork borer, and bacterial growth was quantified.

### Gene expression analysis

Leaf tissues were collected three weeks after TRV inoculation to assess transcript levels in *NbRPF2*-silenced plants. In transgenic plants, expression analysis was conducted 3 weeks post-germination in both *N. benthamiana* and Arabidopsis. Total RNA was extracted using the TRIzol (Sigma) according to the manufacturer’s guidelines. cDNA synthesis was performed using oligo (dT) primers and a reverse transcriptase kit (Thermo Fisher Scientific). RT-qPCR was conducted using SYBR Green dye on an Applied Biosystems Quant 6 machine.

### ABA sensitivity assay and drought stress response of Arabidopsis plants

Seeds of *AtRPF2*-OE, *AtRPF2*-RNAi, and Col-0 were subjected to 1 µM ABA on MS medium, with daily monitoring of germination and calculation of germination rate. Three-week-old Arabidopsis and four-week-old *N. benthamiana* plants, including *NbRPF2-OE*, *NbRPF2-RNAi*, *AtRPL10A*-OE, *AtRPF2*-OE, and RNAi lines, were subjected to drought stress by gradually reducing the water content to 40% of the field capacity over 10 days (Babitha et al., 2013). A control group of plants was maintained at full field capacity. For proteomics, leaf tissues after one week of drought stress were collected. Relative water loss was determined by hourly weighing of detached leaves from Arabidopsis and *N. benthamiana*, with the percentage calculated based on initial leaf weight.

### Stomatal aperture measurement

Leaves from three-weeks-old plants at a similar developmental stage were collected from their adaxial and abaxial leaf surfaces. The epidermal layers of these leaves were observed using a Nikon bright-field microscope at 40x magnification. The number of stomatal openings was measured using ImageJ software (Kaundal et al., 2017).

### Proteomics data generation and analysis

Four-week-old *AtRPF2-OE*, *AtRPL10A-OE,* and respective RNAi lines were infected with the host pathogen and exposed to drought stress. Leaf material was collected for proteomic analysis from each independent stress treatment, three days post-infection, and ten days after drought exposure. One gram of tissue was ground in RIPA buffer (50 mM Tris-HCl, pH 7.4, 1% Triton-X-100, 0.5% DOC, 0.1% SDS, 1mM EDTA, 1mM PMSF) and centrifuged at 13,000 rpm for 30 min at 4 °C. The supernatant was collected, and 500 μg of protein from each sample was precipitated using acetone to remove chlorophyll. The protein pellet was dissolved in 8 M urea. Trypsin (Promega) digestion was performed using 2 M urea-adjusted protein with 100 mM ammonium bicarbonate. For trypsin digestion, 50 μg of protein was used, determined using the PierceTM BCA Protein Assay Kit (REF 23225). For the digestion of 50 μg of protein, 2.5 μg of trypsin was used. Waters OASIS HLB Cartridge C18 was used to purify the peptides before subjecting them to mass spectrometry analysis by Sciex Zenotof 7600.

After acquiring the peptides, MaxQuant_v2.5.2.0 was used for peptide and protein identification. The intensity values obtained from MaxQuant were used to identify differentially regulated proteins using MetaboAnalyst_v6.0 with a Log2 FC ≥ 1.5 and a P-value ≤ 0.05. This list of differentially accumulated proteins was annotated using Uniprot (Proteome ID: UP000006548). Further analysis of these proteins was performed using Venny_v2.1 to identify common and unique proteins, ShinyGo_v0.82 for KEGG pathway and network analysis, and SRplots.

### Hormone quantification

GA and free SA levels were determined using high-pressure liquid chromatography (HPLC) coupled with mass spectrometry (Agilent Technologies) in Arabidopsis and *N. benthamiana* plants, following a streamlined protocol (Almida et al., 2014). Frozen plant tissues (100 mg) were pulverised in liquid nitrogen and extracted using a 70% methanol and 30% water solution. Standard-labelled hormones, specifically 50 pM SA-d6 and GA-d6, were added to the samples while cold, placed on a rotary shaker, and incubated for an hour. The samples were centrifuged at 16,000 × g for 5 min at 4 °C, and the supernatant was air-dried in glass vials. The dried samples were reconstituted in 100 µL of pure methanol, and 5 µL was injected into an Agilent 1290 ultra HPLC system connected to an Agilent 6430 Triple Quad mass spectrometer (Agilent Technologies). The relative quantities of GA and SA were calculated using labelled hormones and expressed as ng per gram of fresh weight (Ramu et al., 2021).

### Protein synthesis measurement

The ability of these plants to synthesise proteins was assessed using a pulse-chase labelling method with [35S]-methionine incorporation. Fifteen-day-old seedlings of Col-0, AtRPF2-OE, RNAi, and wild-type *N. benthamiana*, all 15 days old, were exposed to [35s]-methionine (PerkinElmer) for a duration of six hours on MS media. Subsequently, total protein was extracted, and the labelled protein was quantified using a scintillation counter.

### Yeast two-hybrid (Y2H) assay

Yeast two-hybrid assays were performed using the ProQuest two-hybrid system (Thermo Fisher Scientific). *AtRPF2* was fused to the *GAL4* activation domain in the *pDEST22* prey vector, and *AtRPL10A* was fused to the *GAL4* DNA-binding domain of *pDEST32* as a bait vector. An Arabidopsis cDNA library developed from a mixed elicitor library (Lee et al., 2018) was used to screen for AtRPF2-interacting proteins. Bait and prey constructs were co-transformed in yeast *Mav203* competent cells on synthetic defined medium -Leucine/Tryptophan/Histidine (triple dropout) containing 20 mM 3-aminotriazole (AT). The X-GAL assay was performed using liquid media to confirm the interactions.

### Bi-molecular fluorescence complementation assay

Three-week-old *N. bentamiana* plants were treated with *Agrobacterium* strain GV3101 containing split-GFP vectors *cCFP GFP-AtRPF2* and *nVENUS GFP-AtRPL10A*. GFP fluorescence reconstitution was assessed 48 h later. Observations were made using a Leica SP5 confocal microscope equipped with a 63x oil-immersion lens, using a 405 nm laser for DAPI and a 488 nm laser for GFP. DAPI was added 30 min before image capture.

### Bio-Layer Interferon assay

To study biophysical protein-protein interactions, AtRPF2 was expressed in *pDEST17* with a His-tag vector, whereas AtRPL10 was expressed in *pDEST15* with a GST-tag in *E. coli* Rosetta cells. The proteins were co-incubated with GST resins (Merck), separated by 10% SDS-PAGE, and transferred to a PVDF membrane. AtRPL10A was detected using a GST antibody (Sigma-Aldrich 06-332), and AtRPF2 was identified using a His antibody (Sigma-Aldrich SAB4301134). Proteins purified from *E. coli* were used for biophysical interaction studies using bio-layer interferometry (Octet Red 96). Various concentrations of AtRPF2-His-tag proteins were immobilised onto Ni-NTA biosensors in a buffer containing 50 mM Tris-HCl and 300 mM NaCl. The AtRPL10A-GST tag protein served as the ligand at 50 nM, with subsequent dilutions of 1:3, 1:9, and 1:12. The analysis of biomolecular kinetic interactions involved a loading phase of 500 s, an association phase of 600 s, and a dissociation phase of 600 s. The Octet data analysis software was used to process the raw datasets and calculate the K_on_, K_off_, and Kd values.

### Imaging of Trichomes

To assess unicellular and multicellular trichomes from *N. benthamiana* and Arabidopsis, respectively, leaves from three-week-old plants were collected and placed under vacuum in an environmental scanning electron microscope (Hitachi TM3000) to capture images of the trichomes. For each leaf, over 30 images were captured from at least 10 different biological samples.

### Quantification of glucosinolates

Liquid chromatography–tandem mass spectrometry (LC-MS/MS) was used to quantify glucosinolates in leaf extracts from Arabidopsis lines Col-0, *AtRPF2*-OE, and RNAi/mutants. Arabidopsis leaves were frozen, finely ground in liquid nitrogen, and extracted with 70% methanol, followed by a 15-minute incubation in a hot-water bath. The samples were ultrasonicated for 15 min and then centrifuged at 2,700 × g for 10 min at room temperature. The supernatant was collected, passed through a column, washed three times to eliminate chlorophyll, and extracted using a 20 mM NaOAc buffer for a sulfatase reaction. The NaOAc solution was then injected into the LC-MS/MS system to quantify the untargeted glucosinolates (Hooshmand and Fomsgaard et al., 2021).

### Ribosomal RNA profiling

Total RNA was extracted from four-week-old Arabidopsis plants using TRIzol reagent (Invitrogen) according to the manufacturer’s instructions. Three micrograms of this RNA were run on a 1.2% MOPS/formaldehyde gel (w/v) and then transferred onto a positively charged nylon membrane (Sigma-Aldrich/Merck) using the capillary transfer technique. The RNA probes for 5.8S, 18S, and 25S were 163 and 280 bp, respectively, and were labelled with digoxigenin using the DNA DIG-labelling and detection kit (Roche) according to the manufacturer’s instructions. Hybridisation was performed overnight at 55 °C in a buffer. After hybridisation, the blots were crosslinked with UV light and developed using anti-digoxigenin conjugated to alkaline phosphatase (Sigma). The membrane was then treated with a solution of 5-bromo-4-chloro-3-indolyl phosphate (BCIP) and nitro blue tetrazolium (NBT) to visualise the fragments. Images were captured using a photodetector.

### Statistical analysis

Data from at least three biological datasets were used for the analysis. GraphPad Prism (version 8.0) was used to draw the graphs and perform statistical analyses. Significance values were assessed using the mean values of control or wild-type plants versus those of transgenic treatment plants.

## Author contribution

SY performed the experiments, proteomics, and data analysis; KM performed the northern blotting assay; SS performed BLI and molecular docking; AB helped in proteomics sample preparation and data acquisition; SKR performed the Y2H assay; SD performed MD simulations; TKM helped in proteomics data analysis; and KW generated *N. benthamiana* OE and RNAi lines. KSM and RSV conceptualised the experiments and wrote and edited the manuscript.

## Acknowledgement

This work was supported by the Noble Research Institute, Regional Centre for Biotechnology core grants to RSV; RSV acknowledges a Fulbright-Nehru postdoctoral fellowship from USIEF and a Department of Biotechnology research grant (BT/PR51930/BSA/33/59/2024). SY acknowledges the CSIR Fellowship.

## Data availability

The mass spectrometry proteomics data have been deposited to the ProteomeXchange Consortium via the PRIDE [1] partner repository with the dataset identifier PXD074585

Reviewer access details :

Log in to the PRIDE website using the following details: Project accession: PXD074585

Token: jZo9BlxVXLTN

Alternatively, the reviewer can access the dataset by logging in to the PRIDE website using the following account details: Username, Password: hYTozRcu28aM

---

Table S1: Interacting proteins of AtRPF2 screened through yeast two-hybrid assay.

Supplementary Data S1

Supplementary Data S2

Supplementary Data S3

Supplementary Data S4

Supplementary Data S5

Supplementary Data S6

Supplementary Data S7

Supplementary Data S8

Supplementary Data S9

Supplementary Data S10

Supplementary Data S11

Supplementary Data S12

Figure S1. Phenotype and HR response of *N. benthamiana NbME19A10* silenced plants.

Figure S2: RPF2 is conserved across species and upregulated with different pathogen infections and phenotype of *NbRPF2-*OE and RNAi plants.

Figure S3: Sequence alignment of RPF2 and T-DNA map showing RPF2 mutation.

Figure S4: RNA integrity number and levels of rRNAs in *AtRPF2*-OE, *AtRPF2*-RNAi and mutant lines.

Figure S5: Proteomic changes in *AtRPF2*-OE, *AtRPL10A*-OE, *AtRPF2*-RNAi and *AtRPL10A*-RNAi under normal conditions.

Figure S6: Proteomic changes in *AtRPF2*-OE, *AtRPL10A*-OE, *AtRPF2*-RNAi and *AtRPL10A*-RNAi under pathogen conditions.

Figure S7: Proteomic changes in *AtRPF2*-OE, *AtRPL10A*-OE, *AtRPF2*-RNAi and *AtRPL10A*-RNAi under drought conditions.

