## Supplemental Tables and figures for "Ribosome Processing Factor-2 Interacts with RPL10A to Regulate Selective Translation during Plant Immunity and Drought Stress": Supplementary figures-11.03.2026.pdf

### Supplementary Table and Figures

Table S1: Interacting proteins of RPF2 screened through yeast two hybrid assay.

| | |
|---|---|
| At1g61520 | PSI type III chlorophyll a/b-binding protein |
| At2G16500 | Arginine decarboxylase (ARGdc) |
| AT1G47128 | Cysteine proteinase- RD21a |
| At2g45660 | Suppressor Of Overexpression of Co1 (SOC1) |
| AT1G56010 | No Apical Cotyledon - NAC1 |
| At1g51730 | RWD protein |
| At1g64080 | Membrane-associated kinase regulator 2 |
| AT2G25070 | Protein phosphatase 2C |
| AT5G39190 | Germin type2 protein |
| AT1G08200 | UDP-D-apiose/UDP-D-xylose synthase 2 (AXS2) |
| At2g05100 | Light harvesting complex -B2 |
| AT5g65480 | CLAVATA COMPLEX INTERACTOR |
| AT1G14320 | Ribosomal protein L10 family protein -RPL10 |

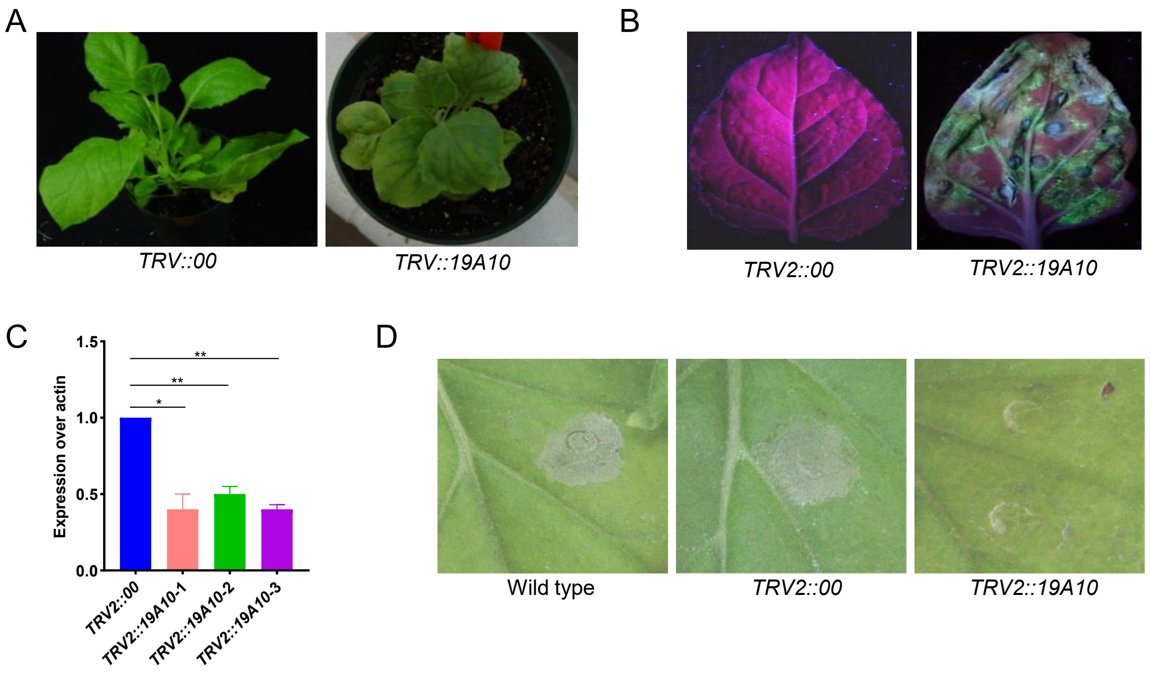


Figure S1. **Phenotype and hypersensitive response (HR) of *N. benthamiana NbME19A10* silenced plants.** A. Dwarf and brittle leaf phenotype of *NbME19A10* silenced plants compared to *TRV::00*. The four-week-old *N. benthamiana* plants were transfected with *NbME19A10* expressed TRV2 and TRV1 constructs along with *TRV::00* constructs. The phenotypic images were taken one month after Agro-infection. B. Bacterial multiplication of nonhost pathogen *P. syringae* pv. *tomato* T1 (*pDSK-dGFPuv*) in *NbME19A10* silenced plants. To the *TRV::19A10* infected plants, after three weeks, nonhost pathogen *P. syringae* pv. *tomato* T1 *(pDSK-dGFPuv*) was vacuum-infiltrated ($1X10^4$ cfu/ml). After 7 dpi, photographs were captured under UV light. C. Transcript levels of *NbME19A10* in VIGS plants. From three-week-old plants, total RNA from *TRV::00* and *TRV::19A10* silenced plants was isolated, and the total transcripts was quantified. *NbActin* is used as a housekeeping gene for normalization. D. Hypersensitive response to nonhost pathogen *P. syringae* pv. *tomato* T1. To silenced and control plants, the nonhost pathogen was syringe-infiltrated, and the lesion area was photographed at 3 dpi.

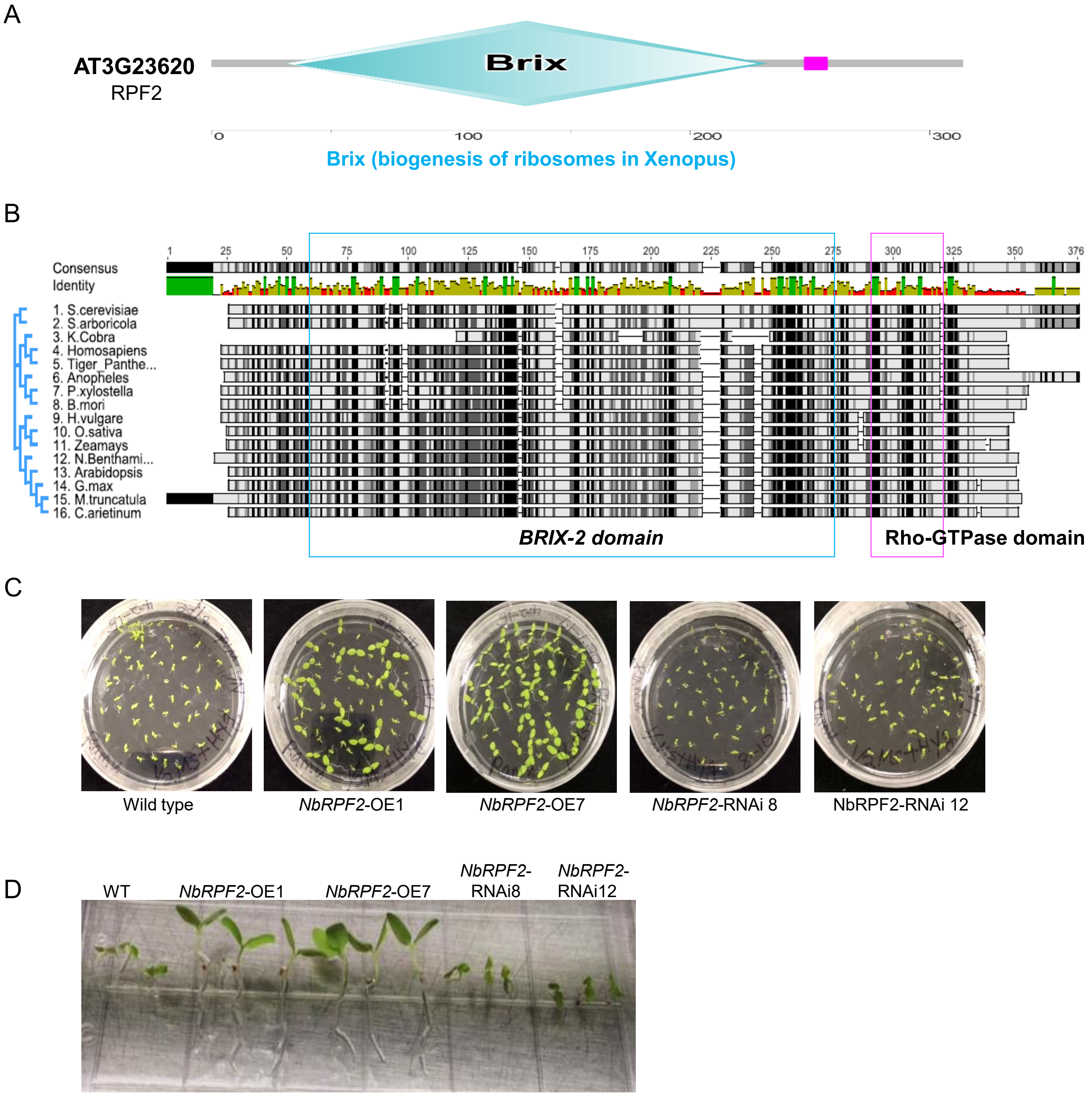


Figure S2: **RPF2 is conserved across species and the phenotype of *NbRPF2*-OE and RNAi plants.** A. *AtRPF2* protein map showing the BRIX domain. The SMART tool was used to predict domain architecture. B. Conserved amino acid sequences of RPF2 in different species showing *BRIX* and GTPase Domain. Geneious bioinformatic tool used to align the amino acid sequences and generate the conserved map. C. *N. benthamiana* seedling germination phenotype expressing *NbRPF2*-OE and RNAi lines. D. Seedling images showing robust root and shoot growth in OE plants compared to RNAi and WT. The full-length *NbRPF2* was cloned and expressed under the 35S-promoter. For RNAi, catalase introns were used and expressed through the Agrobacterium-mediated gene transfer method in *N. benthamaina.* Two independent lines with different transcript levels were used for subsequent characterization.

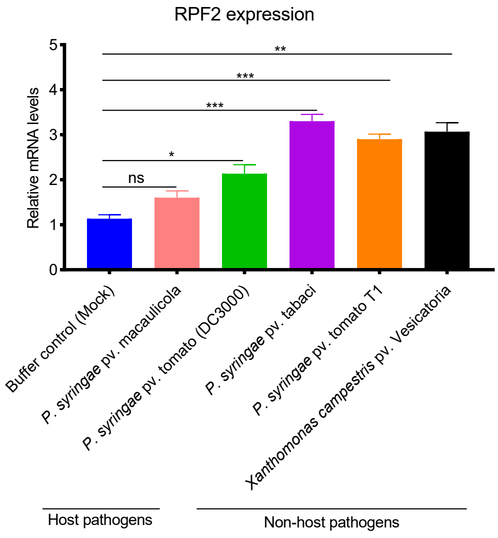


Figure S3. **Expression of *AtRPF2* in Arabidopsis with response to different pathogen.** In-silico expression analysis of *AtRPF2* in Arabidopsis plants infected with pathogens after 3 dpi. The expression data from the public dataset available in TAIR-BAR eFP Browser was obtained from different pathogen-infected samples and histogram was developed.

A Nucleotide

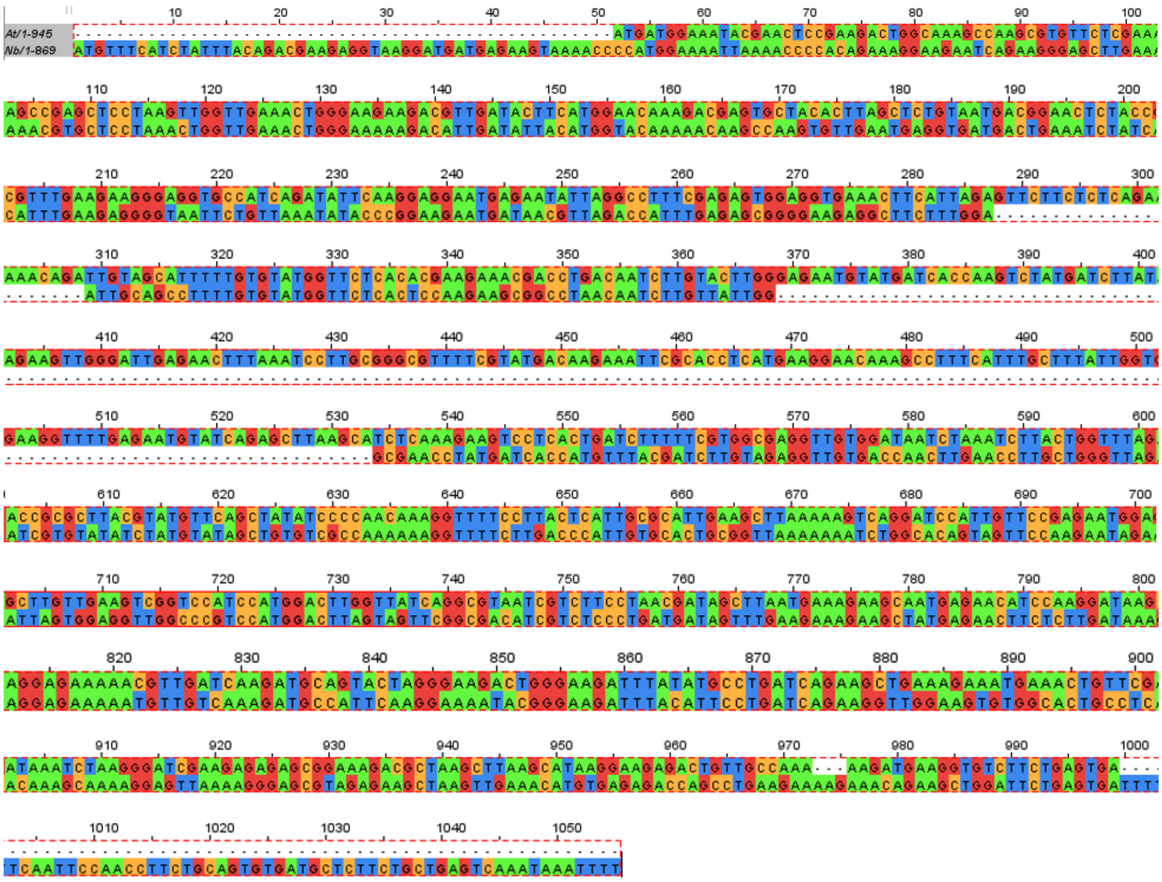

Percent Identity Score- 66.80

B Protein

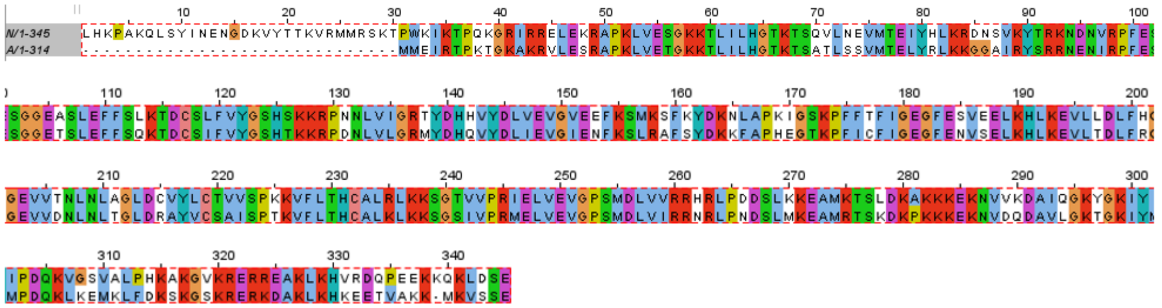

Percent Identity Score- 67.83

C

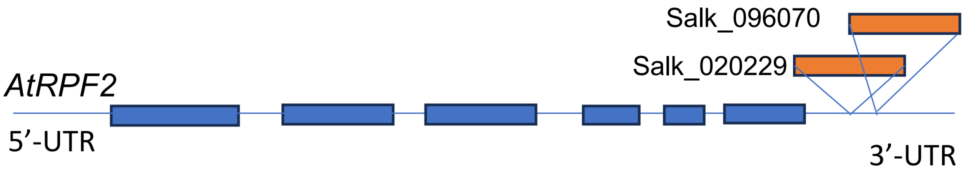


Figure S4: **Sequence alignment of *AtRPF2* with *NbRPF2* and T-DNA insertion sites in the *AtRPF2* gene.** A. Nucleotide sequence alignment showing homology of *NbRPF2* with *AtRPF2*. B. Amino acid sequence showing homology of NbRPF2 and AtRPF2 proteins. C. T-DNA map of Arabidopsis salk-020229 and salk-096070 lines on 3' UTR regions.

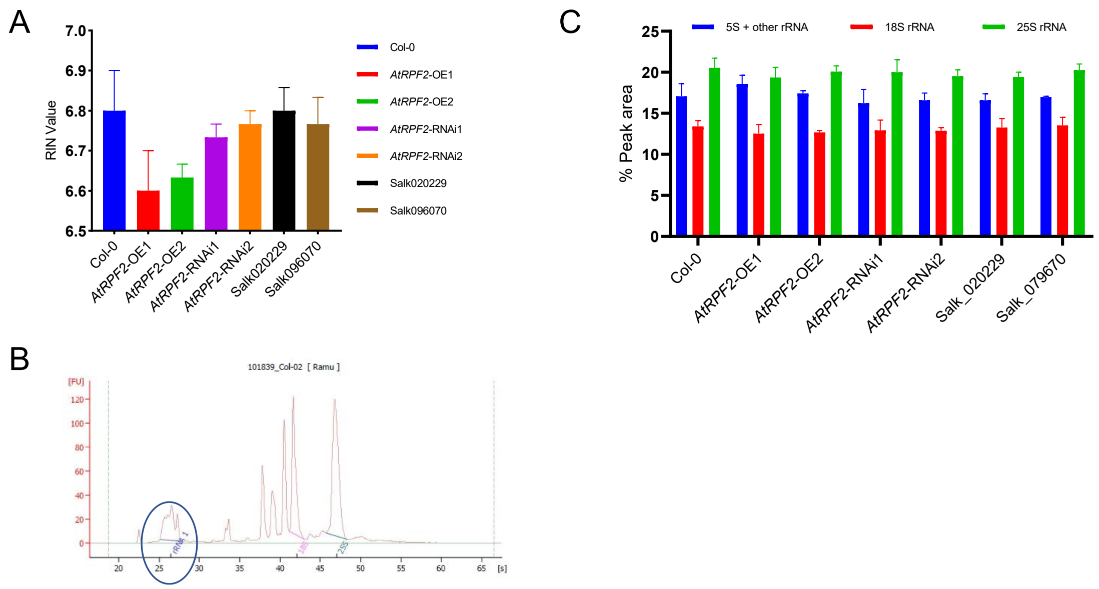


Figure S5: **RNA integrity number and levels of rRNAs in *AtRPF2*-OE, *AtRPF2*-RNAi, and mutant lines.** A. RIN values of total RNA as analysed by BioAnalyzer. B. Chromatogram showing the different peaks of 25S, 18S, and 5.8S rRNA in BioAnalyzer software. C. The levels of 5S rRNA and other rRNA, 18S and 25S rRNA in Col-0, OE, RNAi, and mutant lines. The peak areas from chromatograms were analysed to assess rRNA levels. The error bar indicates replicates from three biological samples.

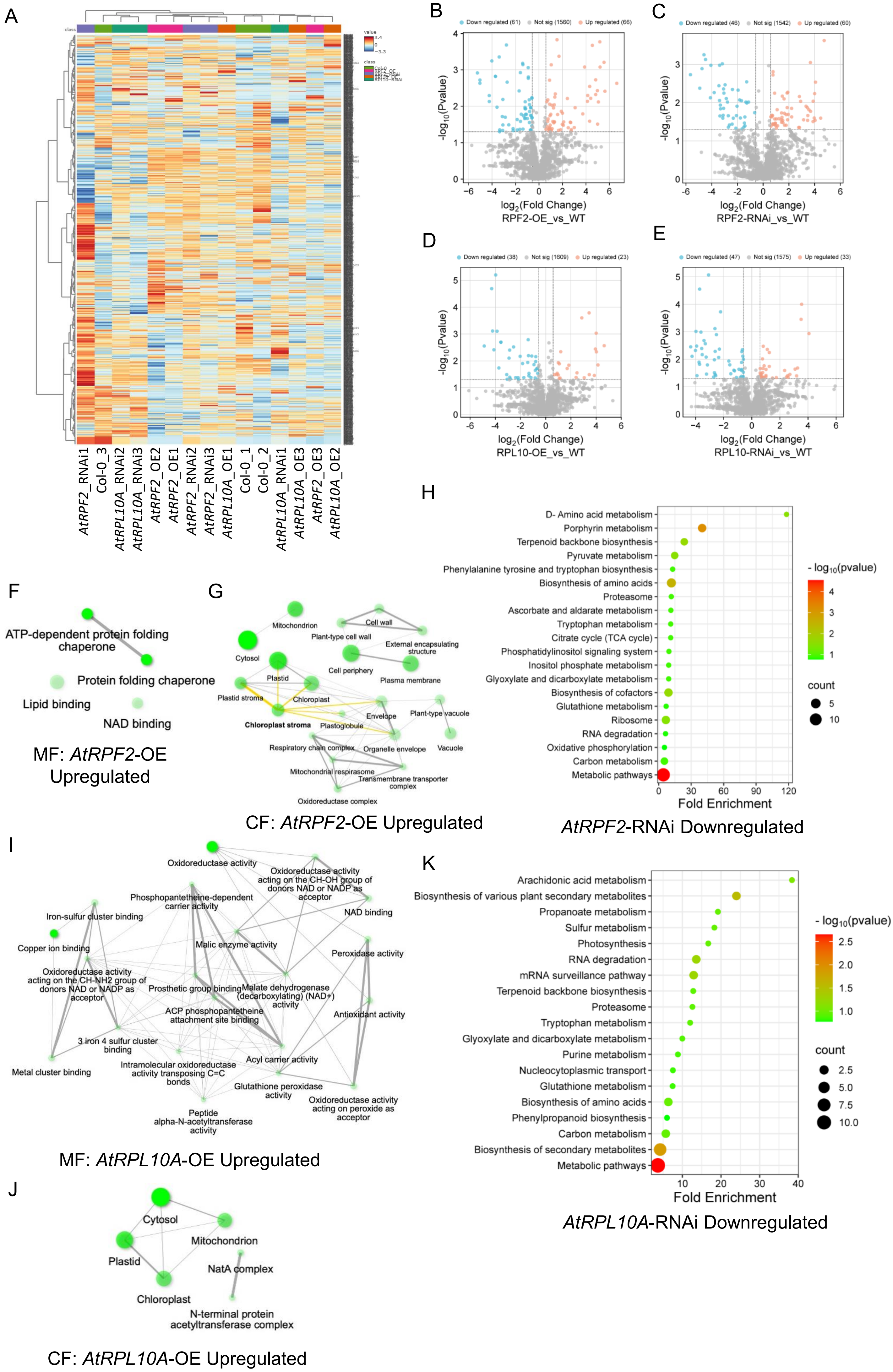
A
AtRPF2_RNAi1
Col-0_3
AtRPL10A_RNAi2
AtRPL10A_RNAi3
AtRPF2_OE2
AtRPF2_OE1
AtRPF2_RNAi2
AtRPF2_RNAi3
AtRPL10A_OE1
Col-0_1
Col-0_2
AtRPL10A_RNAi1
AtRPL10A_OE3
AtRPF2_OE3
AtRPL10A_OE2
B
Down regulated (61) Not sig (1560) Up regulated (66)
-log10(Pvalue)
log2(Fold Change)
RPF2-OE_vs_WT
C
Down regulated (46) Not sig (1542) Up regulated (60)
RPF2-RNAi_vs_WT
D
Down regulated (38) Not sig (1609) Up regulated (23)
RPL10-OE_vs_WT
E
Down regulated (47) Not sig (1575) Up regulated (33)
RPL10-RNAi_vs_WT
F
ATP-dependent protein folding chaperone
Protein folding chaperone
Lipid binding
NAD binding
MF: AtRPF2-OE Upregulated
G
Mitochondrion
Cell wall
Plant-type cell wall
External encapsulating structure
Cytosol
Cell periphery
Plastid
Plasma membrane
Plastid stroma
Chloroplast
Envelope
Plant-type vacuole
Chloroplast stroma
Plastoglobule
Respiratory chain complex
Organelle envelope
Vacuole
Mitochondrial respirasome
Transmembrane transporter complex
Oxidoreductase complex
CF: AtRPF2-OE Upregulated
H
D- Amino acid metabolism
Porphyrin metabolism
Terpenoid backbone biosynthesis
Pyruvate metabolism
Phenylalanine tyrosine and tryptophan biosynthesis
Biosynthesis of amino acids
Proteasome
Ascorbate and aldarate metabolism
Tryptophan metabolism
Citrate cycle (TCA cycle)
Phosphatidylinositol signaling system
Inositol phosphate metabolism
Glyoxylate and dicarboxylate metabolism
Biosynthesis of cofactors
Glutathione metabolism
Ribosome
RNA degradation
Oxidative phosphorylation
Carbon metabolism
Metabolic pathways
Fold Enrichment
- log10(pvalue)
count
AtRPF2-RNAi Downregulated
I
Oxidoreductase activity
Oxidoreductase activity acting on the CH-OH group of donors NAD or NADP as acceptor
Phosphopantetheine-dependent carrier activity
Iron-sulfur cluster binding
NAD binding
Copper ion binding
Malic enzyme activity
Peroxidase activity
Oxidoreductase activity acting on the CH-NH2 group of donors NAD or NADP as acceptor
Prosthetic group binding
Malate dehydrogenase (decarboxylating) (NAD+) activity
ACP phosphopantetheine attachment site binding
Antioxidant activity
3 iron 4 sulfur cluster binding
Acyl carrier activity
Metal cluster binding
Intramolecular oxidoreductase activity transposing C=C bonds
Glutathione peroxidase activity
Oxidoreductase activity acting on peroxide as acceptor
Peptide alpha-N-acetyltransferase activity
MF: AtRPL10A-OE Upregulated
J
Cytosol
Mitochondrion
Plastid
NatA complex
Chloroplast
N-terminal protein acetyltransferase complex
CF: AtRPL10A-OE Upregulated
K
Arachidonic acid metabolism
Biosynthesis of various plant secondary metabolites
Propanoate metabolism
Sulfur metabolism
Photosynthesis
RNA degradation
mRNA surveillance pathway
Terpenoid backbone biosynthesis
Proteasome
Tryptophan metabolism
Glyoxylate and dicarboxylate metabolism
Purine metabolism
Nucleocytoplasmic transport
Glutathione metabolism
Biosynthesis of amino acids
Phenylpropanoid biosynthesis
Carbon metabolism
Biosynthesis of secondary metabolites
Metabolic pathways
Fold Enrichment
- log10(pvalue)
count
AtRPL10A-RNAi Downregulated

Figure S6: **Proteomic changes in *AtRPF2*-OE, *AtRPL10A*-OE, *AtRPF2*-RNAi and *AtRPL10A*-RNAi plants under normal conditions.** A. Differentially expressed proteins from Col-0, OE and RNAi lines of *RPF2* and *RPL10A* are represented in a heatmap. Proteins with more than or equal to 1.5-fold were considered as differentially expressed. Volcano plot analysis showing up- and down-regulated proteins in B. *RPF2*-OE over Col-0 plants. C. *RPF2*-RNAi over Col-0 plants. D. *RPL10A*-OE over Col-0 plants, E. *RPL10A*-RNAi over Col-0 plants. Network of upregulated proteins in *RPF2*-OE plants showing different F. molecular (MF) and G. cellular function (CF). H. Gene set enrichment analysis of downregulated proteins of *RPF2*-RNAi plants. Network showing different hubs of upregulated proteins from *RPL10A*-OE plants showing different I. molecular (MF) and J. cellular function (CF). K. Gene set enrichment analysis of downregulated proteins of *RPL10A*-RNAi showing different KEGG pathways.

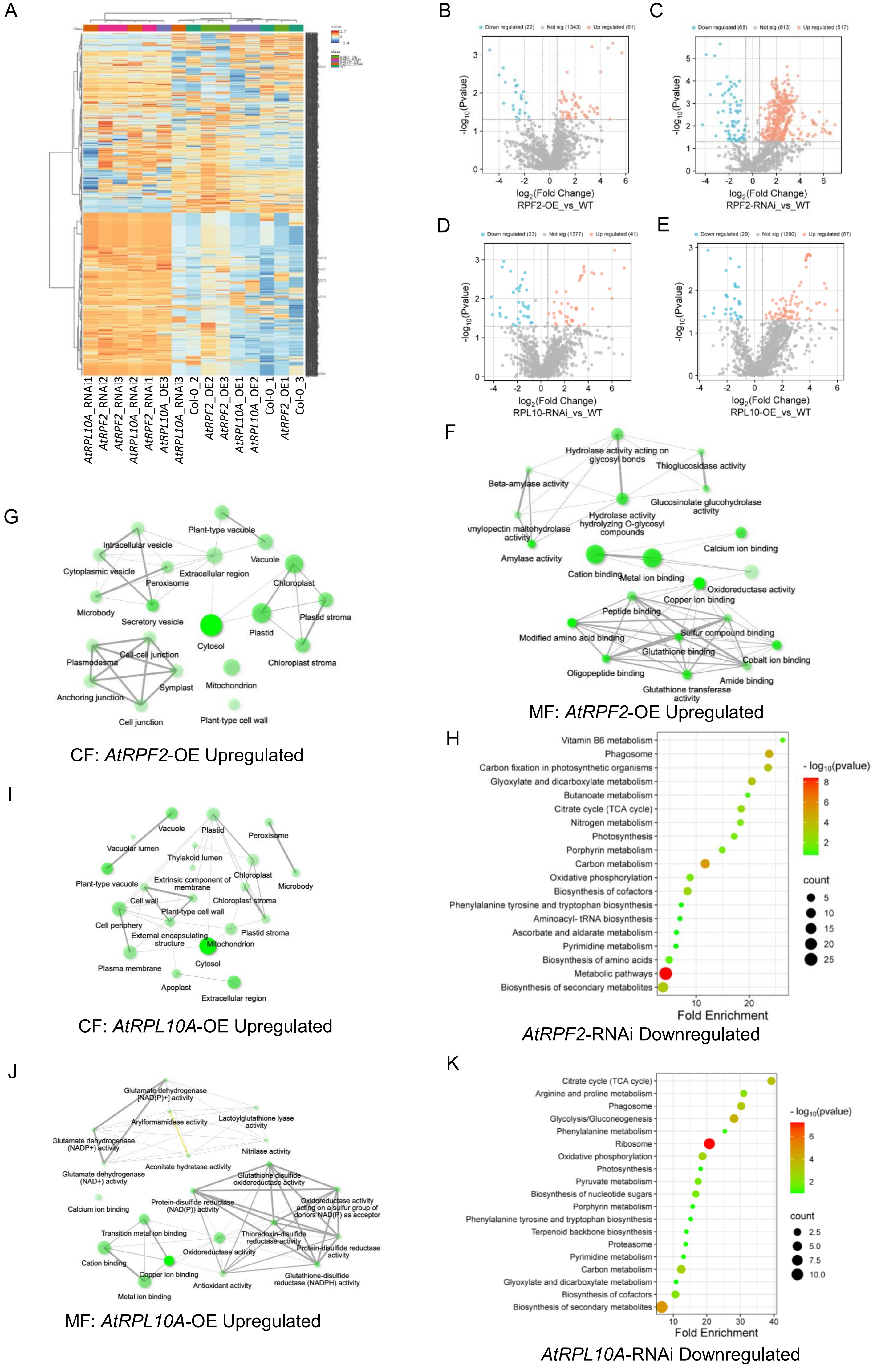

A
AtRPL10A_RNAi1
AtRPF2_RNAi2
AtRPF2_RNAi3
AtRPL10A_RNAi2
AtRPF2_RNAi1
AtRPL10A_OE3
AtRPL10A_RNAi3
Col-0_2
AtRPF2_OE2
AtRPF2_OE3
AtRPL10A_OE1
AtRPL10A_OE2
Col-0_1
AtRPF2_OE1
Col-0_3
B
Down regulated (22)
Not sig (1343)
Up regulated (61)
-log10(Pvalue)
log2(Fold Change)
RPF2-OE_vs_WT
C
Down regulated (68)
Not sig (813)
Up regulated (517)
-log10(Pvalue)
log2(Fold Change)
RPF2-RNAi_vs_WT
D
Down regulated (33)
Not sig (1377)
Up regulated (41)
-log10(Pvalue)
log2(Fold Change)
RPL10-RNAi_vs_WT
E
Down regulated (29)
Not sig (1290)
Up regulated (87)
-log10(Pvalue)
log2(Fold Change)
RPL10-OE_vs_WT
F
Hydrolase activity acting on glycosyl bonds
Thioglucosidase activity
Beta-amylase activity
Glucosinolate glucohydrolase activity
Hydrolase activity hydrolyzing O-glycosyl compounds
Amylopectin maltohydrolase activity
Amylase activity
Calcium ion binding
Cation binding
Metal ion binding
Oxidoreductase activity
Copper ion binding
Peptide binding
Modified amino acid binding
Sulfur compound binding
Glutathione binding
Cobalt ion binding
Oligopeptide binding
Amide binding
Glutathione transferase activity
MF: AtRPF2-OE Upregulated
G
Plant-type vacuole
Intracellular vesicle
Vacuole
Cytoplasmic vesicle
Extracellular region
Peroxisome
Chloroplast
Microbody
Secretory vesicle
Plastid stroma
Plastid
Cytosol
Cell-cell junction
Plasmodesma
Chloroplast stroma
Mitochondrion
Symplast
Anchoring junction
Cell junction
Plant-type cell wall
CF: AtRPF2-OE Upregulated
H
Vitamin B6 metabolism
Phagosome
Carbon fixation in photosynthetic organisms
Glyoxylate and dicarboxylate metabolism
Butanoate metabolism
Citrate cycle (TCA cycle)
Nitrogen metabolism
Photosynthesis
Porphyrin metabolism
Carbon metabolism
Oxidative phosphorylation
Biosynthesis of cofactors
Phenylalanine tyrosine and tryptophan biosynthesis
Aminoacyl- tRNA biosynthesis
Ascorbate and aldarate metabolism
Pyrimidine metabolism
Biosynthesis of amino acids
Metabolic pathways
Biosynthesis of secondary metabolites
- log10(pvalue)
count
Fold Enrichment
AtRPF2-RNAi Downregulated
I
Vacuole
Plastid
Peroxisome
Vacuolar lumen
Thylakoid lumen
Plant-type vacuole
Extrinsic component of membrane
Chloroplast
Microbody
Cell wall
Chloroplast stroma
Plant-type cell wall
Cell periphery
External encapsulating structure
Plastid stroma
Mitochondrion
Cytosol
Plasma membrane
Apoplast
Extracellular region
CF: AtRPL10A-OE Upregulated
J
Glutamate dehydrogenase [NAD(P)+] activity
Arylformamidase activity
Lactoylglutathione lyase activity
Glutamate dehydrogenase (NADP+) activity
Nitrilase activity
Aconitate hydratase activity
Glutamate dehydrogenase (NAD+) activity
Glutathione disulfide oxidoreductase activity
Calcium ion binding
Protein-disulfide reductase (NAD(P)) activity
Oxidoreductase activity acting on a sulfur group of donors NAD(P) as acceptor
Transition metal ion binding
Thioredoxin-disulfide reductase activity
Protein-disulfide reductase activity
Oxidoreductase activity
Cation binding
Copper ion binding
Antioxidant activity
Glutathione-disulfide reductase (NADPH) activity
Metal ion binding
MF: AtRPL10A-OE Upregulated
K
Citrate cycle (TCA cycle)
Arginine and proline metabolism
Phagosome
Glycolysis/Gluconeogenesis
Phenylalanine metabolism
Ribosome
Oxidative phosphorylation
Photosynthesis
Pyruvate metabolism
Biosynthesis of nucleotide sugars
Porphyrin metabolism
Phenylalanine tyrosine and tryptophan biosynthesis
Terpenoid backbone biosynthesis
Proteasome
Pyrimidine metabolism
Carbon metabolism
Glyoxylate and dicarboxylate metabolism
Biosynthesis of cofactors
Biosynthesis of secondary metabolites
- log10(pvalue)
count
Fold Enrichment
AtRPL10A-RNAi Downregulated

Figure S7: **Proteomic changes in *AtRPF2*-OE, *AtRPL10A*-OE, *AtRPF2*-RNAi and *AtRPL10A*-RNAi under pathogen conditions.** A. Heatmap showing differentially expressed proteins from *AtRPF2* and *AtRPL10A,* OE and RNAi lines and Col-0 plants. More than or equal to 1.5-fold were considered as differentially expressed. To the four-week-old Arabidopsis plant, host pathogen *P. syringae* pv*. tomato* (DC3000) was infected and after 3 dpi, the total protein isolated was analyzed using LC/MS-MS. B. Volcano plot analysis showing up- and down-regulated proteins in *AtRPF2*-OE over Col-0. C. *AtRPF2*-RNAi over Col-0. D. *AtRPL10A*-OE over Col-0, E. *AtRPL10A*-RNAi over Col-0. Genetic networks showing different hubs from upregulated proteins in *AtRPF2*-OE plants, showing different- F. molecular (MF) and G. cellular function (CF). H. Gene set enrichment analysis of downregulated proteins from *AtRPF2*-RNAi plants. Upregulated proteins in *AtRPL10A*-OE plants in different networks showing I. molecular (MF) and J. cellular function (CF). K. Gene set enrichment analysis of downregulated proteins from RPL10A-RNAi plants in different KEGG pathways.

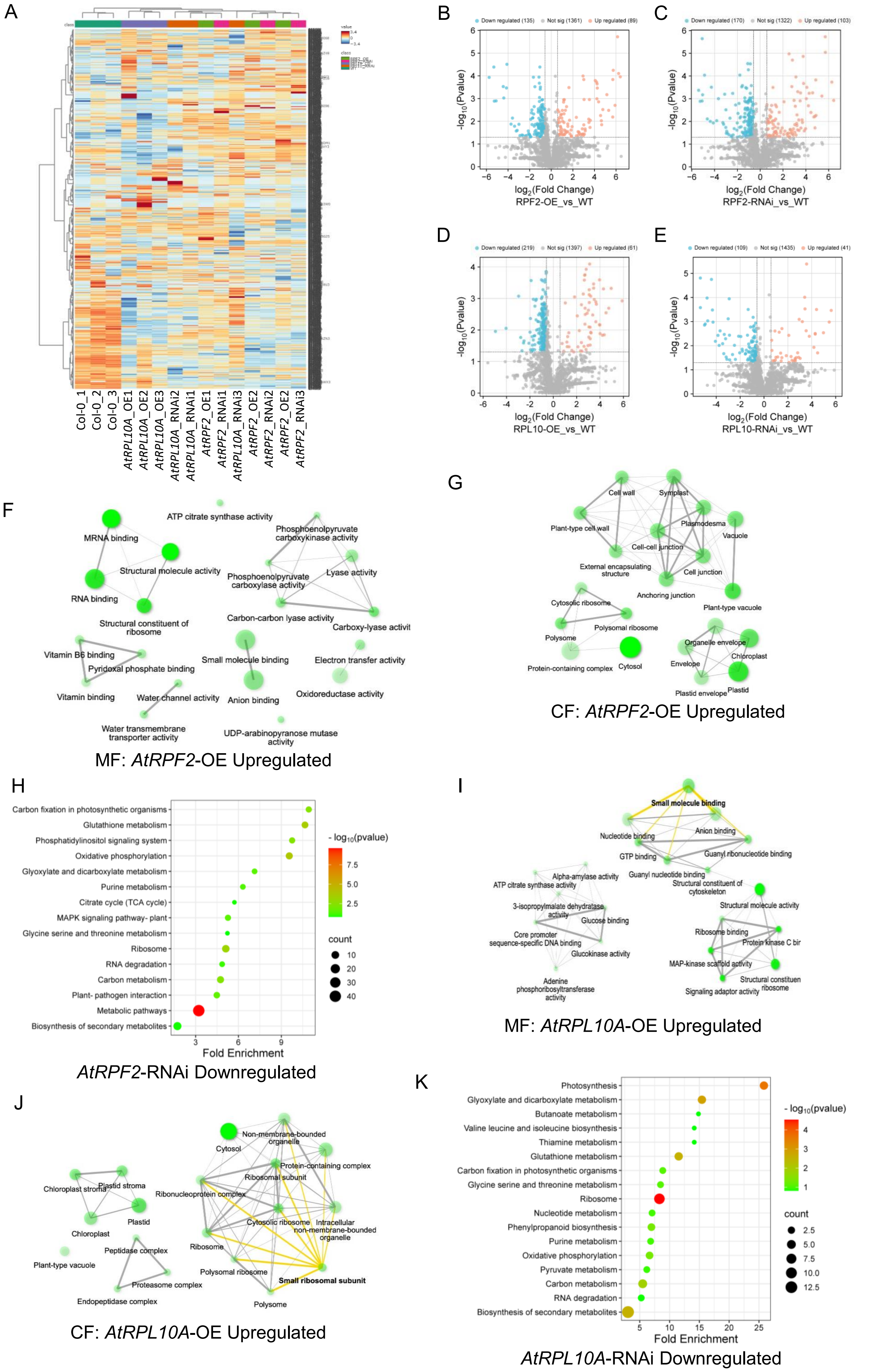
A
Col-0_1
Col-0_2
Col-0_3
AtRPL10A_OE1
AtRPL10A_OE2
AtRPL10A_OE3
AtRPL10A_RNAi2
AtRPL10A_RNAi1
AtRPF2_OE1
AtRPF2_RNAi1
AtRPL10A_RNAi3
AtRPF2_OE2
AtRPF2_RNAi2
AtRPF2_OE2
AtRPF2_RNAi3
B
Down regulated (135)
Not sig (1361)
Up regulated (89)
-log10(Pvalue)
log2(Fold Change)
RPF2-OE_vs_WT
C
Down regulated (170)
Not sig (1322)
Up regulated (103)
-log10(Pvalue)
log2(Fold Change)
RPF2-RNAi_vs_WT
D
Down regulated (219)
Not sig (1397)
Up regulated (61)
-log10(Pvalue)
log2(Fold Change)
RPL10-OE_vs_WT
E
Down regulated (109)
Not sig (1435)
Up regulated (41)
-log10(Pvalue)
log2(Fold Change)
RPL10-RNAi_vs_WT
F
MRNA binding
ATP citrate synthase activity
Phosphoenolpyruvate carboxykinase activity
Structural molecule activity
Lyase activity
Phosphoenolpyruvate carboxylase activity
RNA binding
Structural constituent of ribosome
Carbon-carbon lyase activity
Carboxy-lyase activit
Vitamin B6 binding
Small molecule binding
Electron transfer activity
Pyridoxal phosphate binding
Vitamin binding
Water channel activity
Anion binding
Oxidoreductase activity
Water transmembrane transporter activity
UDP-arabinopyranose mutase activity
MF: AtRPF2-OE Upregulated
G
Cell wall
Symplast
Plant-type cell wall
Plasmodesma
Vacuole
Cell-cell junction
External encapsulating structure
Cell junction
Anchoring junction
Plant-type vacuole
Cytosolic ribosome
Polysomal ribosome
Polysome
Organelle envelope
Chloroplast
Envelope
Protein-containing complex
Cytosol
Plastid envelope
Plastid
CF: AtRPF2-OE Upregulated
H
Carbon fixation in photosynthetic organisms
Glutathione metabolism
Phosphatidylinositol signaling system
Oxidative phosphorylation
Glyoxylate and dicarboxylate metabolism
Purine metabolism
Citrate cycle (TCA cycle)
MAPK signaling pathway- plant
Glycine serine and threonine metabolism
Ribosome
RNA degradation
Carbon metabolism
Plant- pathogen interaction
Metabolic pathways
Biosynthesis of secondary metabolites
Fold Enrichment
- log10(pvalue)
count
AtRPF2-RNAi Downregulated
I
Small molecule binding
Nucleotide binding
Anion binding
GTP binding
Guanyl ribonucleotide binding
Alpha-amylase activity
Guanyl nucleotide binding
ATP citrate synthase activity
Structural constituent of cytoskeleton
3-isopropylmalate dehydratase activity
Glucose binding
Structural molecule activity
Core promoter sequence-specific DNA binding
Ribosome binding
Protein kinase C bir
Glucokinase activity
MAP-kinase scaffold activity
Adenine phosphoribosyltransferase activity
Structural constituen ribosome
Signaling adaptor activity
MF: AtRPL10A-OE Upregulated
J
Cytosol
Non-membrane-bounded organelle
Protein-containing complex
Chloroplast stroma
Plastid stroma
Ribosomal subunit
Ribonucleoprotein complex
Plastid
Chloroplast
Cytosolic ribosome
Intracellular non-membrane-bounded organelle
Peptidase complex
Ribosome
Plant-type vacuole
Polysomal ribosome
Proteasome complex
Small ribosomal subunit
Endopeptidase complex
Polysome
CF: AtRPL10A-OE Upregulated
K
Photosynthesis
Glyoxylate and dicarboxylate metabolism
Butanoate metabolism
Valine leucine and isoleucine biosynthesis
Thiamine metabolism
Glutathione metabolism
Carbon fixation in photosynthetic organisms
Glycine serine and threonine metabolism
Ribosome
Nucleotide metabolism
Phenylpropanoid biosynthesis
Purine metabolism
Oxidative phosphorylation
Pyruvate metabolism
Carbon metabolism
RNA degradation
Biosynthesis of secondary metabolites
Fold Enrichment
- log10(pvalue)
count
AtRPL10A-RNAi Downregulated

Figure S8: **Proteomic changes in *AtRPF2*-OE, *AtRPL10A*-OE, *AtRPF2*-RNAi, and *AtRPL10A*-RNAi under drought conditions.** A. Heatmap showing differentially expressed proteins from Col-0, OE, and RNAi lines of *AtRPF2* and *AtRPL10A*. More than or equal to 1.5-fold was considered differentially expressed. To the four-week-old plants, drought stress was imposed, and tissues were collected after one week. B. Volcano plot analysis showing up- and down-regulated proteins in *AtRPF2*-OE over Col-0. C. *AtRPF2*-RNAi over Col-0. D. *AtRPL10A*-OE over Col-0, E. *AtRPL10A*-RNAi over Col-0 plants. F. Molecular and G. cellular networks from upregulated proteins of *AtRPF2*-OE plants. H. Gene set enrichment analysis of *AtRPF2*-RNAi downregulated proteins. I. Molecular and J. cellular networks from upregulated proteins in *AtRPL10A*-OE plants. K. Gene set enrichment analysis of *AtRPL10A*-RNAi downregulated proteins showing in different KEGG pathways.
